# Fluorogenesis: Inducing Fluorescence in a Non-Fluorescent Protein Through Photoinduced Chromophore Transfer of a Genetically Encoded Chromophore

**DOI:** 10.1101/2023.06.24.546416

**Authors:** Yashwant Kumar, Reman Kumar Singh, Manisha Ojha, Karthik Pushpavanam

**Author notes:** To whom all correspondence must be addressed Karthik Pushpavanam, Ph.D., Discipline of Chemical Engineering, Indian Institute of Technology Gandhinagar, Gujarat, India, 382355, India.

## Abstract

Fluorescent proteins, while essential for bioimaging, are limited to visualizing cellular localization without offering additional functionality. We report for the first time a strategy to expand the chemical, structural, and functional diversity of fluorescent proteins by harnessing light to induce red fluorescence in a previously non-fluorescent protein. We accomplish this by inducing the transfer of the genetically encoded chromophore from a photocleavable protein (PhoCl1) to a non-fluorescent kinase (*Mj*RibK) inducing red fluorescence in the latter. We have employed analytical and spectroscopic techniques to validate the presence of red fluorescence in *Mj*RibK. Furthermore, molecular dynamics simulations were carried out to investigate the amino acid residues of *Mj*RibK involved in the generation of red fluorescence. Finally, we demonstrate the ability of the red fluorescent *Mj*RibK to operate as a cyclable high-temperature sensor. We anticipate that this light-induced chromophore transfer strategy will open new possibilities for developing multifunctional genetically encoded fluorescent sensors.

## INTRODUCTION

Proteins play a vital role in disease emergence, including cancer, making it imperative to comprehensively study their structure, location, and function with precise spatiotemporal resolution.^1^ Among the diverse array of techniques available to visualize and track proteins, fluorescent labeling remains one of the widely adopted techniques.^2^ The most common approach is to genetically append fluorescent proteins to the protein of interest to visualize the complex biological processes.^3^ These fluorescent proteins (∼1000) have been engineered through a combination of experimental and simulation approaches to offer a diverse spectrum of vibrant colors.^4^ Alternatively, the use of genetically encoded peptide tags with probe-specific binding sequences, specific enzyme recognition, and specific protein-protein interactions offer complementary fluorescent methods depending on the intended application.^3^ While these techniques offer remarkable versatility, they lack additional functionalities to quantify and measure concurrent events occurring within the cell. These events encompass a wide range of dynamic processes, including but not limited to pH fluctuations, enzyme activities, protein-protein interactions, and cellular signaling pathways.^5^ Consequently, there is a pressing need to advance the development of genetically encoded fluorescent sensors to effectively capture and measure these concurrent events.

Temperature is a crucial physiological parameter governing the biological status of a living organism.^6^ Cells dynamically interact and adapt to changes in surrounding temperature while simultaneously generating or absorbing heat through exothermic or endothermic reactions during cellular metabolism.^6^ Recognizing the role of temperature in biological processes, significant advancements have been made in the design of genetically encoded fluorescent sensors specifically tailored for temperature monitoring. These sensors operate by detecting temperature variations through changes in fluorescence intensity, enabling precise and reliable temperature measurements. For instance, the green fluorescent protein was used as a tool for mapping intracellular temperature to provide valuable insights into temperature variations within living cells, offering a non-invasive and highly sensitive approach.^7^ A recent study conducted by Lu *et al*. highlights the use of genetically encoded nanothermometers (BgTEMP), consisting of tandem fusions of tdTomato and mNeonGreen fluorescent proteins, to investigate intracellular heat transfer.^8^ Although effective, the existing sensors do not cover a wide range of temperatures, from sub-ambient to hyperthermic conditions. Furthermore, these sensors do not display the ability to recover their initial fluorescence intensity on repeated heating-cooling cycles (cyclability) due to thermally induced destabilization of the fluorescent protein. All these taken together, suggest the pressing need to develop genetically encoded fluorescent temperature sensors that circumvent the above-mentioned limitations.

During our ongoing exploration of fusion constructs with photocleavable fluorescent proteins, we serendipitously discovered a light-induced activation strategy to confer red fluorescence to a previously non-fluorescent protein. The construct involved the fusion of a non-fluorescent riboflavin kinase (*Mj*RibK) from a hyperthermophilic archaeon *Methanocaldococcus jannaschii* to the C-terminus of a photocleavable protein PhoCl1.^9-11^ Surprisingly, when photoexposed, this fusion protein exhibited an intriguing transition from its original green fluorescence to a red fluorescence. This observation is contradictory to the current view where PhoCl1 and its fusion partners loses green fluorescence after exposure to 400 nm light while *Mj*RibK has no known fluorescence emission characteristics.^11-13^ The manuscript describes our efforts in understanding this red fluorogenesis (fluorescence + genesis) through a combined experimental and molecular dynamics simulation approach. We elucidate that the emergence of red fluorescence in *Mj*RibK is due to the site-specific binding of the detached chromophore from PhoCl1 to *Mj*RibK after exposure to 400 nm light. Finally, we demonstrate the potential of this red-fluorescent *Mj*RibK to monitor a wide range of temperatures (25-65 °C) while exhibiting exceptional cyclability. To the best of our knowledge, this unique light activation-mediated generation of red fluorescence in a non-fluorescent protein through a transfer of post-translationally modified genetically encoded chromophores has not been reported elsewhere.

## RESULTS AND DISCUSSION

The fusion construct responsible for the generation of red fluorescence upon exposure to 400 nm light consists of PhoCl1 and *Mj*RibK linked through a KLGGGS linker (**Table S1**). PhoCl1 when exposed to 400 nm light splits into two fragments, (1) an N-terminal fragment (230 amino acids) and (2) a C-terminal fragment (12 amino acids) (**Figure S1**).^11^ The C-terminal fragment possesses the *p*-hydroxybenzylidene-imidazolinone (*p*-HBI) moiety (formed by post-translational modification of HYG residues), followed by an NRVFTKYPR amino acid sequence.^14^ The *p*-HBI moiety confers the characteristic absorption and emission to the native PhoCl1 (λ_Ex_: 380 nm, λ_Em_: 510 nm).^11^ The fusion partner is an archaeal riboflavin kinase (*Mj*RibK) from the hyperthermophile *Methanocaldococcus jannaschii* connected to the C-terminal of PhoCl1 through KLGGGS flexible linker (**Table S1**). For brevity we will refer to the fusion construct as PhoCl1-*Mj*RibK, the C-terminal fragment [*p*-HBI]-NRVFTKYPR-KLGGGS, and the N-terminal empty β-barrel released after photo exposure as [*p*-HBI]-loop and Empty-PhoCl1 barrel respectively.

PhoCl1-*Mj*RibK displays the characteristic green light emission (green fluorescence) of PhoCl1 under the UV-transilluminator (λ_Ex_: 365 nm). We further subjected PhoCl1-*Mj*RibK to 400 nm LED (1.5 mWcm^-2^) for 5 h using our custom-designed light exposure setup (**Figure S2**). Surprisingly, this photo exposure led to an unexpected outcome: the emission of red light (red fluorescence) from the PhoCl1-*Mj*RibK solution. (**Figure 1 A, B**). This observation is in contradiction with the previously reported findings, where an evident decrease in green fluorescence of PhoCl1 and its fusion partners was detected upon exposure to 400 nm light.^11-13^ To validate that the emergence of red fluorescence was specifically attributed to PhoCl1-*Mj*RibK, we conducted control experiments with wild type proteins (PhoCl1 and WT-*Mj*RibK). The unexposed PhoCl1 displays the characteristic green fluorescence while WT-*Mj*RibK does not display any fluorescence under the UV-transilluminator. As anticipated, photoexposure of PhoCl1 led to a decrease in green fluorescence which has been attributed to the detachment of the C-terminal fragment bearing the *p*-HBI from the β-barrel core of PhoCl1 (**Figure 1 A, B, and S1A**).^11-13^ On the contrary, light exposure to WT-*Mj*RibK resulted in no qualitative change in the color of the solution, indicating its fluorescence inactivity. (**Figure 1 A, B**). To validate the requirement of the fusion construct for the generation of red fluorescence, we photoexposed an equimolar amount (20 μM each) of PhoCl1 and WT-*Mj*RibK. The photo-exposed solution did not exhibit red fluorescence highlighting the requirement of a fusion construct. (**Figure 1 A, B**).

**Figure 1.**
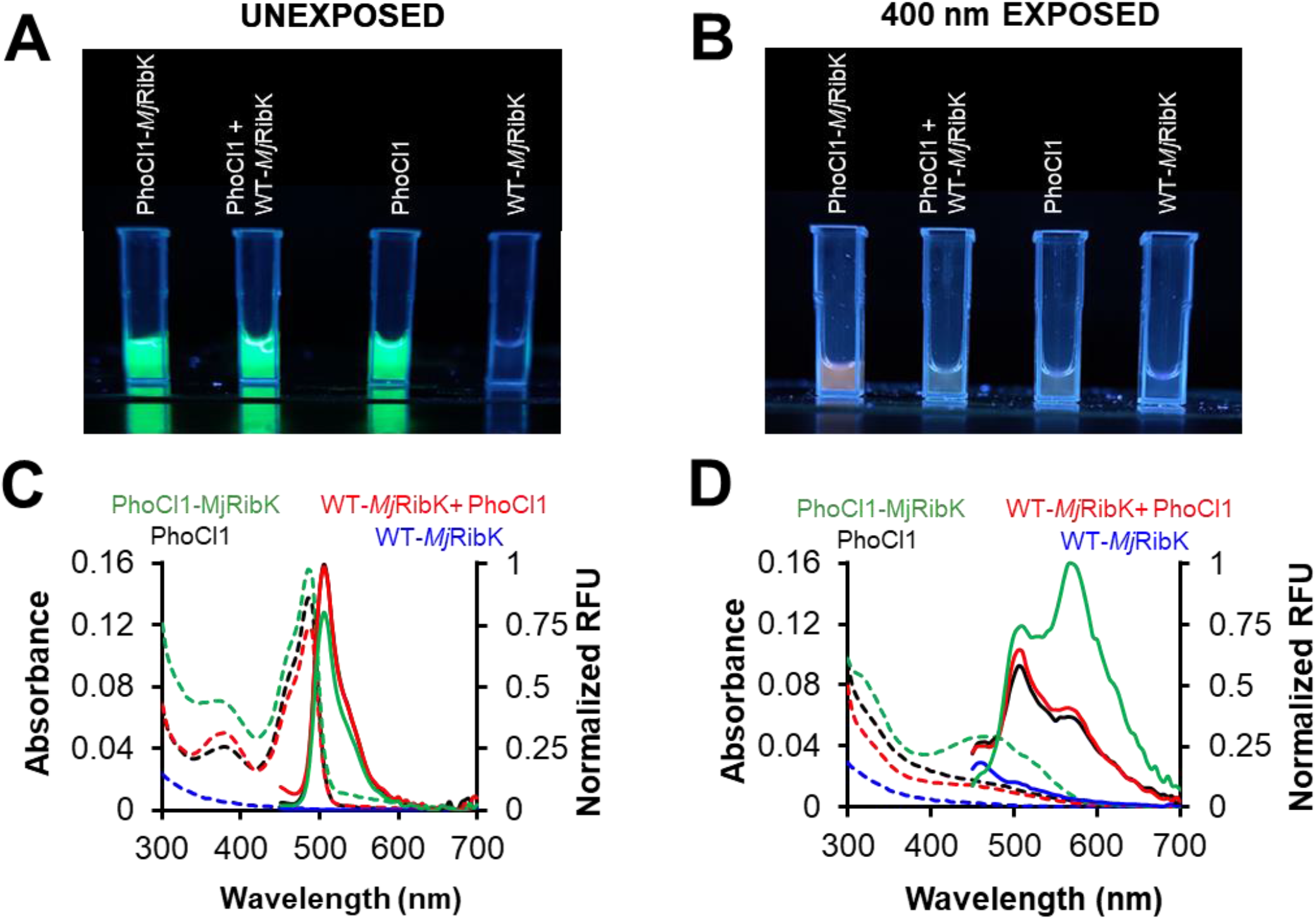
Generation of red fluorescence through photoexposure of PhoCl1-*Mj*RibK **(A)** (From Left to Right) Digital images of PhoCl1-*Mj*RibK, an equimolar mixture of PhoCl1 and WT-*Mj*RibK, PhoCl1, and WT-*Mj*RibK acquired under a UV transilluminator (365 nm) before exposure to 400 nm light. The three samples (Left to Right) enclosed within the cuvette exhibit the characteristic green fluorescence of PhoCl1. The sample (Rightmost) enclosed within the cuvette containing WT-*Mj*RibK is non-fluorescent. **(B)** (From Left to Right) Digital images of PhoCl1-*Mj*RibK, an equimolar mixture of PhoCl1 and WT-*Mj*RibK, PhoCl1, and WT-*Mj*RibK acquired under a UV transilluminator (365 nm) after photoexposure to 400 nm light for 5 h. The PhoCl1-*Mj*RibK exhibits red fluorescence after photoexposure while PhoCl1 and its equimolar mixture with WT-*Mj*RibK exhibit decreased green fluorescence with no visible red emission. WT-*Mj*RibK remains non-fluorescent after photoexposure. **(C)** Absorption-emission spectra of PhoCl1-*Mj*RibK, an equimolar mixture of PhoCl1 and WT-*Mj*RibK, PhoCl1, and WT-*Mj*RibK before exposure to 400 nm light. The dashed and solid lines represent absorption and emission spectra respectively. The samples containing PhoCl1-*Mj*RibK fusion, an equimolar mixture of PhoCl1 and WT-*Mj*RibK and PhoCl1 displayed the characteristic absorption and emission maxima of PhoCl1 at 480 nm and 510 nm respectively. WT-*Mj*RibK displayed no absorption and emission in the range of 300-700 nm (blue solid and dashed lines). **(D)** Absorption-emission spectra of PhoCl1-*Mj*RibK, an equimolar mixture of PhoCl1 and WT-*Mj*RibK, PhoCl1, and WT-*Mj*RibK after exposure to 400 nm light. PhoCl1-*Mj*RibK displayed the characteristic absorption and emission maxima of PhoCl1 at 480 nm and 510 nm respectively. Additionally, the emergence of a new peak centered around 570 nm is indicative of red fluorescence. The absorption and emission spectra of PhoCl1 and its equimolar mixture with WT-*Mj*RibK exhibit a decrease in the absorption and emission maxima of PhoCl1 at 480 nm and 510 nm without the emergence of a new peak centered around 570 nm. WT-*Mj*RibK displayed no absorption and emission in the range of 300-700 nm. These studies utilized a protein concentration of 20 μM.

We characterized the spectral properties of PhoCl1, *Mj*RibK, and PhoCl1-*Mj*RibK using UV-Visible spectroscopy before and after photo exposure (**Figure 1 C, D**). The samples containing PhoCl1-*Mj*RibK, an equimolar mixture of PhoCl1 and WT-*Mj*RibK and PhoCl1 displayed the characteristic absorption and emission maxima of PhoCl1 at 480 nm and 510 nm respectively. WT-*Mj*RibK did not display any characteristic absorption or emission profile between 300 and 700 nm indicating its fluorescence inactivity. Upon photoexposure, a noticeable and expected decrease in the absorption and emission maxima of PhoCl1 and its equimolar mixture with WT-*Mj*RibK was observed. In contrast, the photoexposed PhoCl1-*Mj*RibK displayed a new distinct emission peak centered around 570 nm along with the characteristic emission maxima of PhoCl1 at 510 nm. This further supports the initial visual observation of red fluorescence, confirming the successful generation of red fluorescence through photoexposure of PhoCl1-*Mj*RibK.

To rule out the potential of PhoCl1 exhibiting independent generation of red fluorescence after photo exposure, we subjected PhoCl1 to 405 nm, 535 nm, 635 nm, and 808 nm LASERs (∼13 mWcm^-2^ for 405 nm LASER, ∼65 mWcm^-2^ for others; Time = 1 h). We observed no significant increase in the relative fluorescence units (RFU) at 570 nm (characteristic of red fluorescence) in all LASER conditions (**Figure S3**). These indicate PhoCl1 does not possess the ability to generate red fluorescence independently after photo exposure. Finally, to investigate whether the emergence of red fluorescence was specific to *Mj*RibK or could be achieved with other fusion partners, we generated fusion proteins of PhoCl1 with charged variants of superfolder green fluorescent protein.^15^ The variants (+15)GFP and (−30)GFP vary in the degree of positively (lysine and arginine) and negatively charged (aspartic acid and glutamic acid) amino acid residues.^15^ We photo-exposed these fusion constructs but did not detect qualitative and quantitative changes in the relative fluorescence unit at 570 nm, indicating the absence of red fluorescence (**Figure S4**). All these results taken together underscore the crucial role played by *Mj*RibK and its requirement as a fusion partner to PhoCl1 in achieving the observed red fluorescence after photo-exposure.

To elucidate the origin of red fluorescence after photoexposure of PhoCl1-*Mj*RibK, we employed Ni-NTA chromatography to separate the photocleaved components (Empty-PhoCl1 barrel and [*p*-HBI]-loop-*Mj*RibK). The Empty-PhoCl1 barrel component possesses an N-terminal 6xHis tag, which led us to hypothesize that it could be specifically bound to Ni-NTA resin. In contrast the [*p*-HBI]-loop-*Mj*RibK, having minimal affinity to the resin, could be effectively washed, and subsequently collected for further characterization. As anticipated, the SDS-PAGE gel electrophoresis of the photoexposed solution displayed three bands (lane 2) corresponding to intact PhoCl1-*Mj*RibK (45 kDa; Theoretical molecular weight = 45.034 kDa), Empty-PhoCl1 barrel (28 kDa; Theoretical molecular weight = 27.4 kDa), and [*p*-HBI]-loop-*Mj*RibK (17 kDa; Theoretical molecular weight = 17.7 kDa) (**Figure 2A**). The wash solution displayed a predominant single band (lane 3) corresponding to the molecular weight of [*p*-HBI]-loop-*Mj*RibK. Notably, this solution exhibited red fluorescence as previously observed after photo exposure of PhoCl1-*Mj*RibK.

**Figure 2.**
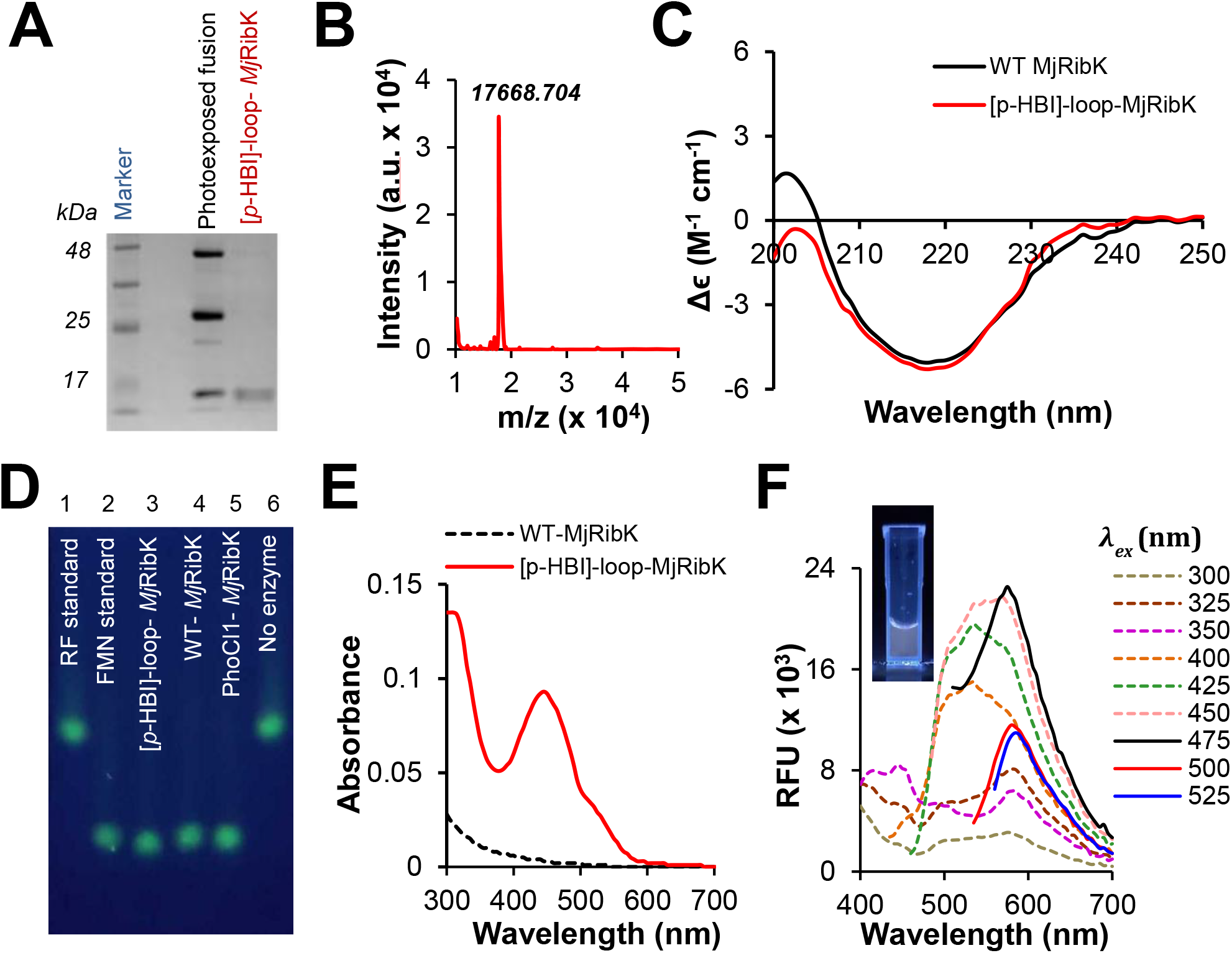
Characterization of [*p*-HBI]-loop-*Mj*RibK. **(A)** 15 % SDS PAGE gel electrophoresis of photoexposed PhoCl1-*Mj*RibK before and after Ni-NTA chromatography. The first lane is the molecular weight marker. The second lane is the photoexposed PhoCl1-*Mj*RibK displaying the bands for PhoCl1-*Mj*RibK (∼ 48 kDa), Empty-PhoCl1 barrel (∼25 kDa) and [*p*-HBI]-loop-*Mj*RibK (∼17 kDa). The third lane represents the wash components after Ni-NTA affinity chromatography. A single band at ∼17 kDa indicates the presence of [*p*-HBI]-loop-*Mj*RibK in the wash solution after Ni-NTA affinity chromatography, **(B)** The MALDI-TOF mass spectrum of the photoexposed components after Ni-NTA affinity chromatography. The spectrum indicates the presence of a single molecular mass peak at 17668.704 Da which is close to the theoretically predicted mass of [*p*-HBI]-loop-*Mj*RibK (17686.79 Da). The MALDI-TOF data is reproduced through the digitization utility of Origin 9.0 software using the original PDF data (**Figure S5**). **(C)** Circular dichroism spectra of WT-*Mj*RibK (black solid line) and [*p*-HBI]-loop-*Mj*RibK (red solid line) indicate minimal changes to the secondary structure **(Table S3). (D)** Fluorescent TLC image displaying the standards and riboflavin kinase activity of the proteins. The first and second lanes correspond to riboflavin (RF) and flavin mononucleotide (FMN) standards respectively. The third and fourth lanes display a single fluorescence spot indicating the conversion of RF into FMN by [*p*-HBI]-loop-*Mj*RibK and WT-*Mj*RibK respectively. The fifth and sixth lanes are the PhoCl1-*Mj*RibK reaction and no enzyme controls. Collectively, the TLC result indicated no difference in the activity between [*p*-HBI]-loop-*Mj*RibK and WT-*Mj*RibK **(E)** UV-Visible absorption spectra of [*p*-HBI]-loop-*Mj*RibK and WT-*Mj*RibK. The [*p*-HBI]-loop-*Mj*RibK shows an absorption maximum between 400-600 nm, whereas WT-*Mj*RibK does not display characteristic absorption. **(F)** Emission spectra of [*p*-HBI]-loop-*Mj*RibK at different excitation wavelengths (300-525 nm). A broad emission spectrum up to excitation wavelength 450 nm is observed while increasing the excitation wavelength to 475 nm yielded a relatively sharp emission peak centered at λ= 580 nm. Further increase in excitation wavelength resulted in a decrease in the emission peak. The inset is a digital image of a cuvette with pure [*p*-HBI]-loop-*Mj*RibK exposed to a UV-transilluminator (365 nm).

We additionally corroborated the purity and molecular weight of the wash solution component by MALDI-TOF analysis. The spectrum highlighted the presence of a protein of mass 17668.704 Da which closely matches the theoretically predicted mass of 17686.79 Da (**Figure 2B, S5**). These provide the support that the component involved in the generation of the red fluorescence is the photocleaved C-terminal component [*p*-HBI]-loop-*Mj*RibK.

We performed circular dichroism (CD) spectroscopy to validate whether the generation of red fluorescence is attributed to changes in the secondary structure of [*p*-HBI]-loop-*Mj*RibK, (**Figure 2C**). The comparison of the major secondary structure elements (15.7 % helix, and 68.5 % β-sheets) of [*p*-HBI]-loop-*Mj*RibK are in close agreement with those of WT-*Mj*RibK (18.7 % helix, and 65.7 % β-sheets) (**Table S3**). Additionally, we validated the retainment of the three-dimensional structure through a riboflavin kinase activity assay. The assay involves the phosphorylation of riboflavin (RF) into flavin mononucleotide (FMN) using cytidine-5’-triphosphate as a phosphate donor, by the enzymatic action of *Mj*RibK (**Figure S6A**).^9^ By leveraging the distinct migration rates of RF (Lane 1, retention factor = 0.5) and FMN (Lane 2, retention factor = 0.2), we successfully separated these compounds on a thin-layer chromatography (TLC) plate using the upper layer of solvent-mixture 1-butanol, acetic acid and water in 4:1:5 ratio (**Figure 2D**).^16^ The intrinsic fluorescence of riboflavin and FMN allowed us to visualize them on a UV transilluminator. Notably, the enzymatic reactions with [*p*-HBI]-loop-*Mj*RibK (Lane 3), WT-*Mj*RibK (Lane 4), and PhoCl1-*Mj*RibK (Lane 5) enzymes displayed a single spot on the TLC plate at a retention factor of 0.2, indicating complete conversion of RF to FMN. We quantified the fluorescence intensity of the FMN spot on the TLC plate for the [*p*-HBI]-loop-*Mj*RibK (Lane 3) and WT-*Mj*RibK (Lane 4), reactions, revealing no significant variations in enzyme activity between [*p*-HBI]-loop-*Mj*RibK, and WT-*Mj*RibK, while the no enzyme control (Lane 6) displayed minimal fluorescence intensity at the corresponding FMN spot. (**Figure S6B**). These taken together indicate overall structural similarity between [*p*-HBI]-loop-*Mj*RibK and the WT-*Mj*RibK.

We characterized the spectral characteristics of [*p*-HBI]-loop-*Mj*RibK through UV-Visible and fluorescence spectroscopy. The absorption spectrum of [*p*-HBI]-loop-*Mj*RibK displayed a maximum at 450 nm, accompanied by an additional shoulder centered around 525 nm (**Figure 2E**). On the contrary, WT-*Mj*RibK does not display any characteristic absorption spectra between 300-700 nm. This suggests the role of the [*p*-HBI]-loop moiety from PhoCl1 imparting the absorption characteristics to the previously non-chromogenic WT-*Mj*RibK. We recorded the emission spectra of [*p*-HBI]-loop-*Mj*RibK across various excitation wavelengths within its absorption range (**Figure 2F**). When excited with 300 nm, 325 nm, and 350 nm, the fluorescence spectra of [*p*-HBI]-loop-*Mj*RibK exhibited a broad spectrum with two prominent peaks of weak intensities: one at approximately 500 nm and another at around 580 nm. The observed emission spectra are characteristic of photoconvertible red proteins, including those derived from mMaple, such as PhoCl1.^11^ The peak emission intensity increased when the excitation wavelength was increased within the range of 400-450 nm while preserving the broad profile. The broad spectra exhibited an asymmetrical nature, suggesting the presence of three major electronic transitions with emission peaks at approximately 500 nm, 535 nm, and 570 nm. A further increase in excitation to 475 nm resulted in a sharp emission peak with a red shift to 580 nm. Finally, a decrease in the emission peak was observed with increasing the excitation wavelength above 475 nm. We quantified the fluorescence lifetime of [*p*-HBI]-loop-*Mj*RibK as 1.2 ns which is comparable to red fluorescent proteins (mCardinal = 1.3 ns) and Turbo650 = 1.5 ns) (**Figure S7**).^17^ These spectroscopic investigations validate the successful generation of red fluorescence in [*p*-HBI]-loop-*Mj*RibK and highlight the potential of [*p*-HBI]-loop-*Mj*RibK as a red fluorescent protein with a suitable fluorescence lifetime for potential applications in fluorescence-based sensing.

We adopted a molecular dynamics approach to elucidate the mechanism underlying the emergence of red fluorescence in [*p*-HBI]-loop-*Mj*RibK. We focused on identifying the specific binding site of the *p*-HBI chromophore to the Loop-*Mj*RibK. However, *Mj*RibK could feature multiple binding sites for the *p*-HBI chromophore. Therefore, to explore potential binding sites, we generated three initial conformations where the N-terminal *p*-HBI chromophore was positioned distal to the protein core by a distance greater than 25 Å (**Figure S8**). Subsequently, we conducted three separate 1 μs simulations for each conformation and collected an extensive range of conformations from these trajectories which were utilized for further analysis.

We analyzed various *p*-HBI binding sites produced from these simulations by evaluating the total energy (consisting of electrostatic and van der Waals interactions) between the *p*-HBI and the residues present on the loop (referred to as E1), as well as between the *p-*HBI and *Mj*RibK (referred to as E2) for all conformations. The distribution of E1 and E2 provides insight into the interaction energetics of *p*-HBI with the loop and *Mj*RibK. A high E1 and low E2 are indicative of a strong interaction between *p*-HBI and the loop respectively. Conversely, a low E1 and high E2, suggest a strong interaction between *p*-HBI and *Mj*RibK respectively. When E1 and E2 are comparable, it signifies a simultaneous interaction of *p*-HBI with both the loop and *Mj*RibK. A two-dimensional histogram of E1 and E2 revealed the presence of three distinct populations denoted as C1, C2, and C3 (**Figure 3A**). The most representative conformations of [*p*-HBI]-loop-*Mj*RibK at C1, C2, and C3 states are indicated in **Figure 3 B, S9A, and S9B** respectively.

**Figure 3.**
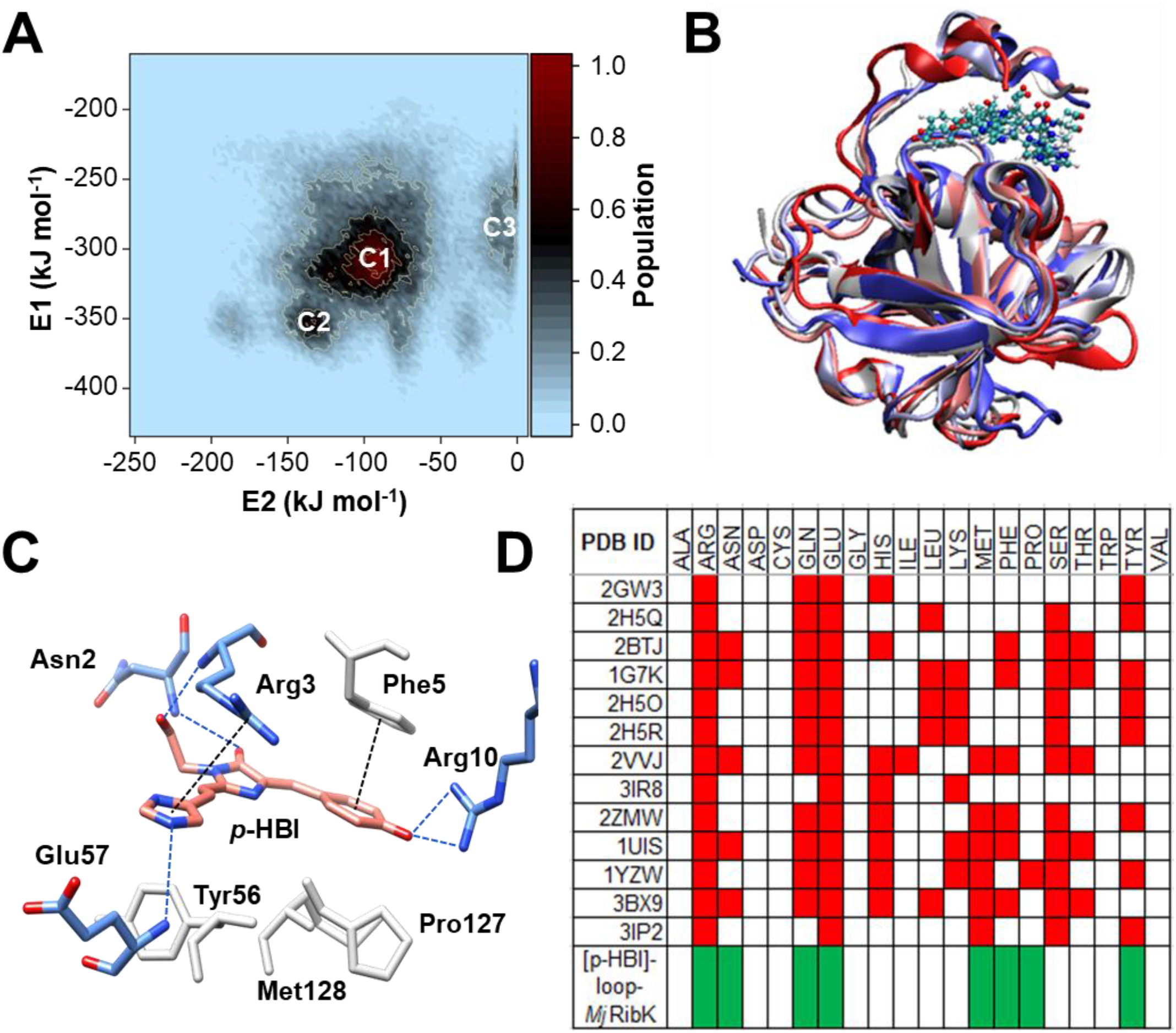
Molecular dynamics simulations of [*p*-HBI]-loop-*Mj*RibK **(A)** Two-dimensional histogram of energy of *p*-HBI with loop (E1) and *p*-HBI with *Mj*RibK (E2). The contours represent the population of the corresponding state. The letter ‘C1’, ‘C2’ and ‘C3’ represent the three most populous states of the system. The population is normalized with the highest populated state (C1). A color gradient is utilized to depict the population distribution, with maroon denoting the most populous state and light blue indicating the least populated one. **(B)** Representation of 50 conformations of *p*-HBI-loop-*Mj*RibK at C1 state. All the conformations are superimposed to show the structural deviation. The protein is shown in ribbon representation and *p*-HBI is shown in ball-stick representation. Each color signifies a distinct protein conformation. These conformations indicate that there is negligible alteration in conformation between all the C1 states. **(C)** The residues of the loop and *Mj*RibK interacting with *p*-HBI chromophore. The residues Asn2, Arg3, Phe5, Arg10 are the residues from the loop itself and are involved in the shielding of *p*-HBI chromophore from water molecules. The residues Glu57, Tyr56, Met128, and Pro127 are from the *Mj*RibK surface and are involved in the interaction with *p*-HBI. The blue dotted lines represent the hydrogen bonds, the black dotted lines represent π-π stacking interactions, and the residues shown in the gray-white are hydrophobic residues. (**D)** List of amino acids near the chromophore in different red fluorescent proteins. The red cell represents the presence of the amino acid residue near the chromophore of the corresponding protein (Column 1). The green cell represents the presence of the amino acid residue near *p*-HBI in the [*p*-HBI]-loop-*Mj*RibK at C1 state. This highlights a significant overlap between the key amino acid residues of the [*p*-HBI]-loop-*Mj*RibK near *p*-HBI and amino acid residues near the chromophores of existing red fluorescent proteins.

C1 represents the most populous state of [*p*-HBI]-loop-*Mj*RibK followed by C2 and C3 states. Interestingly, this occurs despite the C2 state exhibiting higher overall energy stability (−350 kJ mol^-1^ and E2 of -140 kJ mol^-1^) compared to C1 (E1 is -300 kJ mol^-1^ and E2 is -90 kJ mol^-1^) and C3 (E1 of -10 kJ mol^-1^ and E2 of - 280 kJ mol^-1^). This observation can be attributed to enthalpy-entropy compensation often observed in protein-drug complexes.^18-20^ For instance, the binding of carbonic anhydrase II to acetazolamide is driven by a favorable enthalpy change, but results in a decrease in system entropy.^21, 22^ Similarly, Williams and co-workers, investigated numerous protein-ligand complexes using isothermal titration calorimetry (ITC) measurements. The statistical analyses revealed that a more favorable enthalpy change was often accompanied by a compensatory decrease in system entropy or *vice-versa* during protein-ligand interaction.^20^ Analogously, we hypothesize an enthalpy-entropy compensation phenomenon suggesting that while the C2 state may exhibit higher energy stability, it likely experiences a compensatory decrease in entropy. In contrast, the C1 state, despite having lower energy stability, could benefit from a more favorable entropy change. This highlights the potential interplay between enthalpy and entropy in the binding of *p*-HBI to *Mj*RibK, where different energetic contributions can offset each other to achieve overall stability and population distribution.

Further, to identify the probable conformation of [*p*-HBI]-loop-*Mj*RibK, we employed a clustering approach to the group structures near the C1, C2, and C3 states (**Figure 3 B, S9A, and S9B**). We utilized a root mean square deviation cutoff of 1 Å, to successfully cluster ∼5000 conformations at C1 state. The 50-representative conformation, superimposed on each other, at C1, C2, and C3 states are shown in **Figure 3 B**, and **S9** respectively. Since the C1 state is the most populous state, we hypothesize that it is the representative of the experimental conformation of [*p*-HBI]-loop-*Mj*RibK in solution and is used for further analysis. We calculated the total interaction energy between the *p*-HBI chromophore and the surrounding amino acids to gain insights into the environment of the *p*-HBI in its C1 state (**Figure S10**). The amino acids Asn2 and Arg3 displayed the strongest interaction energies (∼ -100 kJ mol^-1^) with *p*-HBI. The residues phenylalanine (Phe5), glutamic acid (Glu57), tyrosine (Tyr56), proline (Pro127), glutamine (Gln133), and methionine (Met128) exhibited weaker interaction energies of ∼ -20 kJ mol^-1^. The residues Glu57, Arg3, and Arg10 are involved in hydrogen bonding with the imidazole nitrogen while the residues Tyr56, Pro127, and Met128 (shown in gray-white) contribute to hydrophobic interactions. Additionally, we observe π-π stacking interactions of the *p*-HBI with Arg3 and Phe5. These interactions facilitate the orientation of *p*-HBI in the vicinity of *Mj*RibK (**Figure 3C**). We compared these amino acid residues to the amino acid residues that are commonly present around the chromophore in red fluorescent proteins from the literature (**Figure 3D**).^23-32^ We observed that among the 20 natural amino acids, there is a high degree of conservation of Arg, Asn, Gln, Glu, His, Leu, Lys, Met, Phe, Ser, Tyr, and Thr. Notably, there is a remarkable similarity between the amino acids observed in our study and those reported to surround the chromophore required for red fluorescence. We postulate that the red fluorescence is generated due to the proximity of the *p*-HBI to these amino acid residues - Asn2, Arg3, Phe5, Glu57, Tyr56, Pro127, and Met128.

The fluorescence emission by a fluorescent protein is highly dependent on the planar conformation of the chromophore within the interior β-barrel fluorescent protein. To validate whether the *p*-HBI satisfies this criterion, we measured the dihedral between the C1-C2-C3-C4 atoms in *p*-HBI (**Figure S11**). This revealed a population predominantly ranging between 171°-179° suggesting a nearly planar orientation of *p*-HBI in [*p*-HBI]-loop-*Mj*RibK (**Figure S11**). Additionally, the presence of a hydrophobic environment is critical for facilitating light emission by limiting fluorescence loss through non-radiative processes.^33-36^ To verify the generation of a hydrophobic environment, we calculated the total number of water molecules around the *p*-HBI chromophore during its transition from an unbound state to bound state (C1) by identifying all water molecules within a 4 Å distance cut-off. We observe a clear decrease in the total number of water molecules from ∼50 to ∼10 when transitioning from the unbound to the bound state (**Figure S12**). This observation indicates the binding of *p*-HBI to the *Mj*RibK surface involves the displacement of initially bound water molecules. These molecular features of planar conformation and the decrease in the total number of water molecules near the *p*-HBI chromophore in [*p*-HBI]-loop-*Mj*RibK maximize the efficiency of light emission while minimizing fluorescence quenching through non-radiative pathways. Collectively, these simulations offer insights into the potential molecular mechanisms underlying the red emission displayed by [*p*-HBI]-loop-*Mj*RibK following 400 nm photoexposure to PhoCl1-*Mj*RibK.

Genetically encoded fluorescent temperature sensors have emerged as valuable tools to investigate thermal fluctuations in biological systems, ranging from subcellular compartments to whole organisms.^7, 8^ Although various proteins have been developed, these suffer from an inability to (1) Operate at high temperatures (> 50 °C) and/or (2) Cyclability (heating-cooling cycle). Our previous results indicate the role of the [*p*-HBI]-loop in the red fluorogenesis of [*p*-HBI]-loop-*Mj*RibK. We hypothesize that with increasing temperature, the [*p*-HBI]-loop within the protein will undergo conformational changes along with displacement from the amino acid residues involved in the generation of red-fluorescence. These combined effects would ultimately lead to a loss in fluorescence. Conversely, we anticipate fluorescence restoration with a decrease in temperature through the acquisition of the initial C1 state. We suggest leveraging this temperature-dependent fluorescence change to create a genetically encoded fluorescent sensor for cyclic monitoring of temperature. Furthermore, the exceptional heat tolerance of *Mj*RibK (up to 100 °C), will likely allow for reliable high-temperature sensing where conventional proteins might not be suitable due to their limited thermal tolerance.^9, 37^ Indeed, we observe a linear decrease (R^2^ ∼0.99) in fluorescence with increasing temperatures up to 65 °C (**Figure 4A**). We also observe the recovery of fluorescence with decreasing temperature back to its original temperature of 25 °C. The [*p*-HBI]-loop-*Mj*RibK demonstrates complete recovery of the fluorescence even after 5 thermal cycles between 25-65 °C (**Figure 4B**).

**Figure 4.**
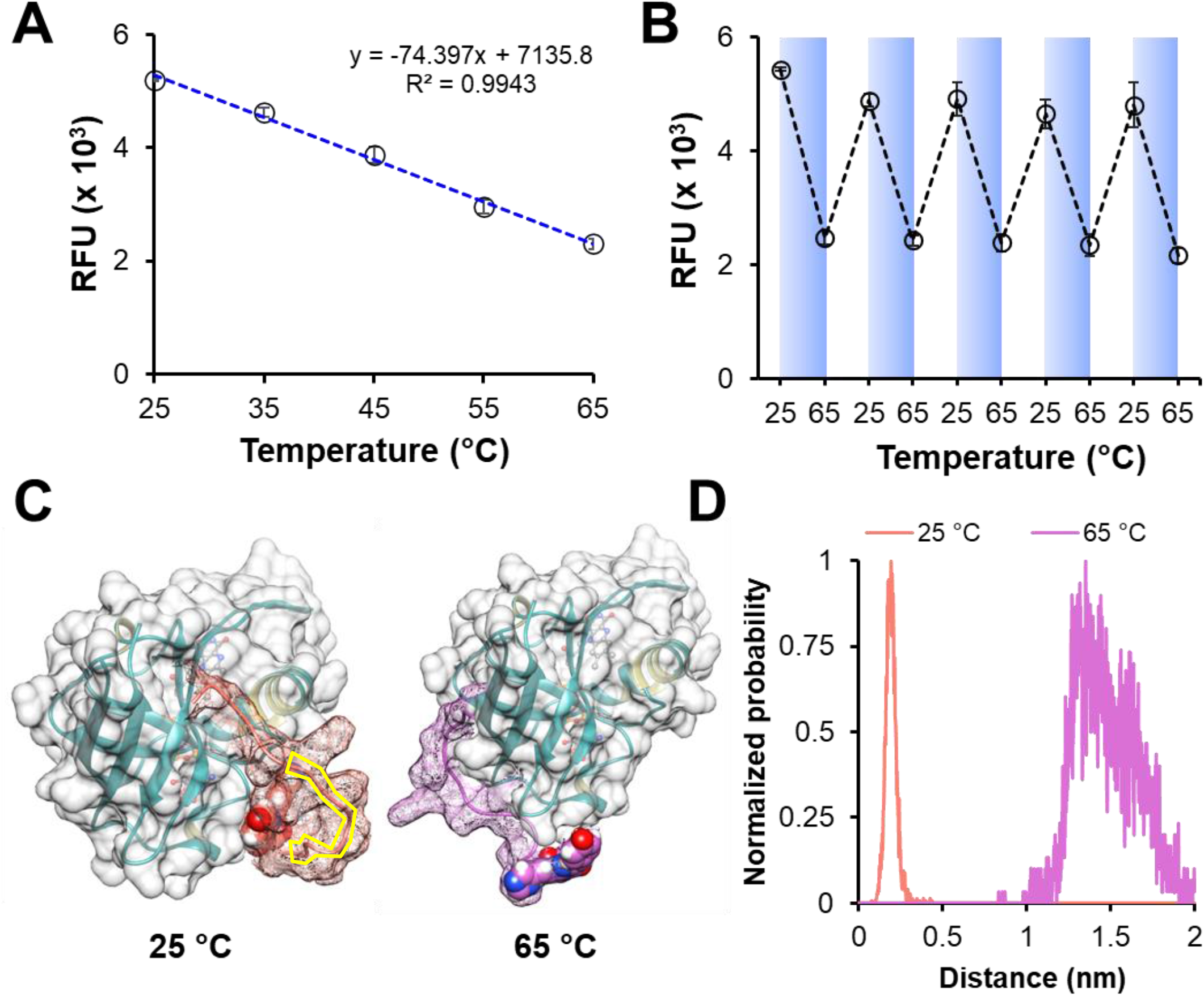
Effect of temperature on the emission characteristics of [*p*-HBI]-loop-*Mj*RibK **(A)** A plot of relative fluorescence units (RFU) as a function of temperatures indicates a linear decrease in RFU of [*p*-HBI]-loop-*Mj*RibK with an increase in temperature from 25 °C to 65 °C, **(B)** A plot of change in RFU of [*p*-HBI]-loop-*Mj*RibK over 5 cycles of heating and cooling between 25 °C and 65 °C. [*p*-HBI]-loop-*Mj*RibK regains its RFU after cooling to 25 °C from 65 °C, **(C)** The energy minimized PDB structures of [*p-*HBI]-loop-*Mj*RibK at 25 °C (Left) and 65 °C (Right). The [*p*-HBI]-loop segment at 25 °C and 65 ° are highlighted with salmon-red and orchid-purple mesh surfaces respectively. A part of the loop (highlighted inside the yellow box) forms a protective outer layer, effectively shielding the *p*-HBI from water molecules. However, at 65 °C, *p*-HBI and the entire connecting loop (highlighted in orchid-purple) adopt an extended conformation and simultaneously exposing the *p*-HBI to the water molecules. **(D)** Change in center of mass distance between *p-*HBI and *Mj*RibK at 25 °C (salmon-red) and 65 °C (orchid-purple). An increase in the distance by 0.5-1 nm with increasing the temperature from 25 °C to 65 °C likely contributes to the observed loss in fluorescence with increasing temperature.

We corroborated our hypothesis by performing temperature-dependent molecular dynamics simulations of [*p-*HBI]-loop-*Mj*RibK at 25 °C and 65 °C. **Figure 4C** illustrates the energy-minimized structures of [*p*-HBI]-loop-*Mj*RibK at 25 °C (left) and 65 °C (right), respectively. At 25 °C, a segment of the loop (highlighted within the yellow box) forms a protective outer layer, effectively shielding the *p*-HBI chromophore from water molecules. However, at 65 °C, *p*-HBI and the entire connecting loop (highlighted in orchid-purple) adopts an extended conformation. Notably, this extension of the *p*-HBI conformation at 65 °C is accompanied by an increase in the number of water molecules (∼35) surrounding the chromophore (**Figure S12**). We also calculated the distance between the center of masses (COM) of *p*-HBI and the protein at 25 °C and 65 °C (**Figure 4D**). Our analysis provides support for the displacement of *p*-HBI from the surface of *Mj*RibK, with a noticeable increase in distance (>1 nm) at 65 °C. In contrast, at 25 °C, the *p*-HBI remains in much closer proximity to the protein, with a distance of less than 0.5 nm. The increase in distance between the COM of *p*-HBI and the protein and the increase in the total number of water molecules surrounding *p*-HBI likely contribute to the observed loss in fluorescence with increasing temperature. This provides compelling evidence for the temperature-dependent behavior of [*p*-HBI]-loop-*Mj*RibK but also underscores its potential as a genetically encoded fluorescent sensor for monitoring temperature changes. The ability of this system to withstand high temperatures and exhibit cyclic behavior positions it as a highly promising candidate for applications demanding precise and repeatable temperature sensing in various biological contexts.

## CONCLUSION

To the best of our knowledge, we have successfully demonstrated the first light-induced strategy to induce red fluorescence in a previously non-fluorescent protein through the transfer of a genetically encoded chromophore. Our strategy involved the creation of a fusion protein between a fluorescent photocleavable protein (PhoCl1) and a non-fluorescent riboflavin kinase (WT-*Mj*RibK). Through systematic experimentation, we validated the emergence of red fluorescence in WT-*Mj*RibK was indeed the result of light-induced chromophore transfer from PhoCl1. We quantified the identity and integrity of the now-transformed red fluorescent *Mj*RibK through SDS-PAGE and MALDI-TOF spectroscopy. The experimental measurements closely aligned with the theoretically calculated values, validating the binding of the genetically encoded chromophore to *Mj*RibK. In addition, we employed circular dichroism spectroscopy and an enzyme activity assay to evaluate whether the emergence of red fluorescence in *Mj*RibK resulted from alterations in its structural composition. Our results indicated no substantial changes in the protein’s structure when compared to the WT-*Mj*RibK, further supporting the role of the genetically encoded chromophore in the induction of red fluorescence. We conducted molecular dynamics simulations, to gain insights into the mechanisms underlying the generation of red fluorescence. These simulations identified three distinct potential conformations of *Mj*RibK in conjunction with the chromophore, with one conformation exhibiting a significant population. Further analysis of this conformation unveiled the key amino acid residues near the chromophore, which closely resembled those found in existing red fluorescent proteins. Furthermore, we showcased the translational potential of the red fluorescent *Mj*RibK by demonstrating its ability to monitor temperatures as high as 65 °C. We also assessed its cyclability by subjecting it to five cycles of repeated heating and cooling between 25 °C and 65 °C. In each cycle, the red fluorescent protein demonstrated consistent recovery of its fluorescence intensity, reaffirming its reliable performance. Collectively, our results offer exciting opportunities for expanding the chemical, structural, and functional design space of genetically encoded fluorescent proteins with additional functionalities. By utilizing light-induced chromophore transfer, we anticipate new possibilities for developing versatile and multifunctional fluorescent sensors that can be employed in a wide range of biological applications.

## Supporting information

SI_Fluorogenesis: Inducing Fluorescence in a Non-Fluorescent Protein Through Photoinduced Chromophore Transfer of a Genetically Encoded Chromophore

## AUTHOR CONTRIBUTIONS

K.P conceived the original idea and planned the experiments. Y.K carried out the experiments and data analysis. R.K.S conceived, planned and analyzed the simulation data. M.O assisted Y.K during experiments. K.P supervised the research along with providing feedback during the writing of the manuscript. All authors provided critical feedback and helped shape the research, analysis, and manuscript.

## ACKNOWLEDGEMENTS

The authors thank CRTDH (Common Resource and Technology Development Hub), Cell Culture lab, and CIF (Central Instrumentation Facility), Nanoplasmonics lab (Professor Saumyakanti Khatua, Dr. NVS Praneeth and Dr. Diptiranjan Paital) and Proteomics and Peptide Synthesis facility (PPSF) for providing various instrumentation facilities at IIT Gandhinagar. We acknowledge Krishna Agrawal, Akshant Kumawat and Gitanjali Swarup for assisting in experiments. The authors would also like to acknowledge National supercomputing facility Param Brahma, (IISER Pune) and IIT Bombay supercomputing facility for providing the necessary computational facilities. We acknowledge the generous support of Gujarat State Biotechnology Mission and Startup Research Grant IIT Gandhinagar provided to K.P.

