## Supplementary material for "Fluorogenesis: Inducing Fluorescence in a Non-Fluorescent Protein Through Photoinduced Chromophore Transfer of a Genetically Encoded Chromophore": SI_Fluorogenesis: Inducing Fluorescence in a Non-Fluorescent Protein Through Photoinduced Chromophore Transfer of a Genetically Encoded Chromophore

*To whom all correspondence must be addressed

Karthik Pushpavanam, Ph.D.

Discipline of Chemical Engineering

Indian Institute of Technology Gandhinagar

Gujarat, India, 382355, India

**MATERIALS AND METHODS**

**Kits and Materials.** Plasmid purification kits and PCR purification kits, Ni-NTA agarose were purchased from Qiagen. Q5® Hot Start High-fidelity 2X master mix, NEBuilder® HiFi DNA assembly master mix, NcoI-HF® and XhoI restriction enzymes were acquired from New England Biolabs (NEB).

**Chemicals and Reagents.** Tris base, 2-mercaptoethanol, kanamycin sulfate, nickel sulfate, riboflavin, flavin mononucleotide (FMN), sodium chloride, 1-butanol, acetic acid, methanol, glycerol, and, bacterial growth media- Luria Bertani (LB) broth (Miller), LB agar (Miller), were purchased from Sisco Research Laboratory (SRL) Pvt. Ltd. Cytidine-5’-triphosphate (CTP) was procured from Cayman Chemical (USA). Phenylmethylsulfonyl fluoride (PMSF) and disodium dihydrogen ethylenediaminetetraacetate dihydrate (EDTA) was acquired from Tokyo Chemical Industry (TCI) Co. Ltd. Isopropyl-ß-D-1-thiogalactopyranoside (IPTG), low EEO agarose, and imidazole, were obtained from HiMedia Laboratories.

**Consumables.** Thin layer chromatography (TLC) plates and 10 kDa MWCO Amicon® centrifugal filters were acquired from Merck. SnakeSkin^TM^ dialysis tubing (10 kDa MWCO, 0.22 mm Internal diameter) was acquired from Thermofisher Scientific. UV LEDs (400-405 nm, VAOL-5GUV0T4) were purchased from VCC Optoelectronics. Costar® 96-well UV-transparent microplates and Costar® fluorescence black microplates were purchased from Corning.

**Plasmids.** *Mj*RibK plasmid was a generous gift from Dr. Amrita Hazra (Whytamin Lab IISER Pune (India). pET-PhoCl1-6xHis was a gift from Robert Campbell (Addgene plasmid #164033; http://n2t.net/addgene:164033; RRID: Addgene_164033). pET-6xHis-(-30)GFP was a gift from David Liu (Addgene plasmid #62936; http://n2t.net/addgene:62936; RRID: Addgene_62936). pET-6xHis-(pos15)GFP was a gift from David Liu (Addgene plasmid #89248; http://n2t.net/addgene:89248; RRID:Addgene_89248)

**Cloning of Fusion Constructs.** We engineered three fusion proteins by genetically linking the photocleavable protein PhoCl1 to the N-terminal end of (1) A Riboflavin kinase (*Mj*RibK) from *Methanocaldococcus jannaschii*, (2) Super-negatively charged protein ((-30)GFP) and (3) super-positively charged protein ((+15)GFP) using a flexible linker KLGGGS. The pET28a(+) vector was linearized using the NcoI-HF and XhoI restriction enzymes. The gene sequences of PhoCl1, *Mj*RibK, (-30)GFP, and (+15)GFP were PCR amplified from the pET-PhoCl1-6His, pET28a-*Mj*RibK, pET-6xHis-(-30)GFP and pET-6xHis-(pos15)GFP plasmids to include the overhangs required for further processing. The NEB Q5®-Hot Start high-fidelity 2X master mix was used for PCR amplification, following the manufacturer's instructions. The linearized vector and the amplified gene sequences were combined in a 1:2 ratio with the NEB HiFi DNA assembly master mix according to the company's protocol and incubated at 50 °C for 15 min to facilitate DNA assembly. Subsequently, the DNA assembly was transformed into chemically competent *E. coli* DH5α cells. Plasmids were obtained from single colonies that grew after transformation and their sequences were verified using Sanger sequencing (Eurofins Genomics, India). The plasmid containing the fusion gene sequence was then transformed into *E. coli* BL21(DE3) expression cells and used for the overexpression and purification of the PhoCl1 fusion protein. For brevity, the fusion constructs will be referred to as PhoCl1-*Mj*RibK, PhoCl1-(-30)GFP and PhoCl1-(+15)GFP respectively.

**Tables S1 and S2** list the nucleotide/amino acid sequences and primers used for the cloning in this study respectively.

**Overexpression of PhoCl1-*Mj*RibK fusion and other proteins.** *Escherichia coli* BL21(DE3) expression cells, transformed with the plasmids- pET-6xHis-PhoCl1-*Mj*RibK, pET-PhoCl1-6xHis, pET-6xHis-*Mj*RibK, pET-6xHis-PhoCl1-(-30)GFP and pET-6xHis-PhoCl1-(+15)GFP were used to inoculate a 5 mL culture of Luria-Bertani (LB) broth supplemented with 50 µg/mL of kanamycin sulfate (antibiotic). To initiate the culture, a 1% overnight growth inoculum was introduced into 500 mL of LB media supplemented with 50 µg/mL of kanamycin sulfate (antibiotic). The resulting secondary culture was then incubated at 37 °C and 150 rpm in an incubator shaker. Once the optical density at 600 nm of the culture reached 0.6, induction was performed by adding 1 mM IPTG to trigger the overexpression of 6xHis-PhoCl1-*Mj*RibK, PhoCl1, wild type (WT) *Mj*RibK, 6xHis-PhoCl1-(-30)GFP and 6xHis-PhoCl1-(+15)GFP. Following induction, the temperature was reduced to 18 °C and maintained for 18 h. Subsequently, the cells were harvested by centrifugation at 6,000 rpm at 4 °C for 10 mins. The resulting cell pellets were flash-frozen and stored at -80 °C for subsequent utilization.

**Purification of PhoCl1-*Mj*RibK fusion and other proteins.** The cells were resuspended in a lysis buffer consisting of 100 mM Tris-HCl pH 8.0, 300 mM NaCl, 20 mM imidazole, 0.1 mM PMSF, and 0.025 % 2-mercaptoethanol. Ultrasonication was utilized for cell disruption, using a cycle of 1 s on and 3 s off at 60 % amplification for a total time of 20 min. After sonication, the lysed cells were centrifuged at 14,000 rpm for 30 min to separate the cell debris from the supernatant. The clarified lysate was filtered through a 0.22 µm syringe-driven filter and loaded onto a pre-equilibrated 2 mL Ni-NTA gravity column. The equilibration buffer (20 mM imidazole buffer) comprised 20 mM imidazole, 100 mM Tris-HCl pH 8.0, 300 mM NaCl, 0.1 mM PMSF, and 0.025 % 2-mercaptoethanol for all proteins except the supercharged fusion proteins. The purification of supercharged fusion proteins was done using 2 M of NaCl in the buffers instead of 300 mM NaCl. The column was washed successively with 30 mL of equilibration buffer and 30 mM imidazole buffer. Finally, the protein was eluted using a 100 mM imidazole buffer. Ni-NTA affinity purification was also employed to purify the WT-*Mj*RibK. Wash buffers, containing increasing concentrations of imidazole buffer (20 mM, 40 mM, and 70 mM), were utilized to remove non-specific proteins. The elution of WT-*Mj*RibK was achieved using a 250 mM imidazole buffer. The eluted fractions containing the pure fusion proteins were verified through SDS-PAGE gel electrophoresis on a 12 % gel, while those of PhoCl1, WT-*Mj*RibK, and its variant were verified on 15 % SDS-PAGE gel. The pure protein fractions were combined and dialyzed using SnakeSkin^TM^ dialysis tubing against 20 mM Tris-HCl pH 8.0, and 300 mM NaCl.

**Photoexposure Setup.** The positive and negative terminals of the 400 nm LED were placed in separate columns of a breadboard. A 470 kΩ resistor was connected in series with the positive terminal of the 400 nm LED. The negative terminal of the LED and the positive terminal of the resistor was connected to a DC power supply source. The schematic of the circuit and the complete setup is shown in **Figure S2 A, B.** The LED was illuminated by applying 5 V with a set DC of 0.20 mA.

**Photoexposure of Proteins.** To facilitate photoexposure of PhoCl1, WT-*Mj*RibK, PhoCl1-*Mj*RibK, PhoCl1-(-30)GFP and PhoCl1-(+15)GFP, the breadboard-mounted LED was placed in an inverted position inside the microcentrifuge tube, ensuring its proximity to the protein sample **(Figure S2 C)**.

300 µL of the corresponding aqueous protein solution (20 µM) was placed in a 2 mL microcentrifuge tube. The protein solutions were exposed to 400 nm LED light (power: 1.5 mWcm^-^²). The photoexposure was carried out for 5 h for PhoCl1-*Mj*RibK fusion and 8 h for supercharged fusion proteins. As a control, a separate PhoCl1-*Mj*RibK fusion protein sample with the same volume and concentration was shielded from light by covering the tube with aluminum foil. After photoexposure, the change in the colors of protein solutions in comparison to the controls were visualized under a UV transilluminator.

**Purification of [*p*-HBI]-loop-*Mj*RibK.** We aliquoted 900 µL of the purified 6xHis-PhoCl1-*Mj*RibK fusion protein into three individual 2 mL microcentrifuge tubes, with each tube containing 300 µL of the protein solution. Subsequently, these tubes were subjected to a 5 h exposure to 400 nm light with an intensity of 1.5 mWcm^-2^ for the photocleavage of 6xHis-PhoCl1-*Mj*RibK fusion protein. Following light exposure, we loaded the protein solution onto a pre-equilibrated Ni-NTA gravity column. The equilibration buffer contained 20 mM Tris-HCl and 300 mM NaCl at pH 8.0. The N-terminal 6xHis-tagged PhoCl1 barrel was effectively captured by the Ni-NTA resin, while the flow-through fraction contained the photocleaved *Mj*RibK protein. The molecular weight of the purified component was assayed by 15 % SDS-PAGE gel electrophoresis and MALDI-TOF spectroscopy.

**Matrix-Assisted Laser Desorption/Ionization Time-of-Flight Mass Spectrometry (MALDI-TOF MS).** The purified protein solutions were dialyzed against 50 mM (NH4)_2_CO_3_ buffer containing 0.025% 2-mercaptoethanol before MALDI-TOF analysis. A saturated solution of sinapinic acid (3,5-dimethoxy-4-hydroxycinnamic acid) in a mixture of acetonitrile and 1 % TFA (1:2 ratio) was used as the matrix. The materials used for the preparation of the matrix came with the Bruker starter kit for MALDI-TOF MS (Part # 8208241). We mixed equal volumes of matrix and protein samples and drop-casted them on the MALDI target plate and allowed them to dry at room temperature. The MALDI-TOF spectra were recorded on the Bruker Autoflex MALDI-TOF instrument in the linear positive mode from m/z 10000-60000.

**Enzyme Activity Assay of WT-*Mj*RibK and [*p*-HBI]-loop-*Mj*RibK.** The enzymatic activity of WT-*Mj*RibK and [*p*-HBI]-loop-*Mj*RibK was monitored through a riboflavin kinase activity assay.^1^^,^ ^2^ To a mixture of 200 µM riboflavin, 50 mM Tris-HCl pH 8.0, 10 mM cytidine-5′-triphosphate (CTP), 1 mM MgCl_2_, 5 µM of either WT-*Mj*RibK or [*p*-HBI]-loop-*Mj*RibK was added. The samples were incubated at 70 °C for 4 h. The conversion of riboflavin into FMN was estimated through thin-layer chromatography (TLC). 1 µL reaction mixtures along with the standards and controls were spotted on a TLC plate. The mobile phase used to run the TLC was the upper layer of the mixture of solvents- 2-butanol, acetic acid, and water in a 4:1:5 ratio.^3^ The fluorescent spots on the TLC plate corresponding to the reactant ‘riboflavin’ and the product ‘FMN’ were captured on a UV transilluminator using long UV light (365 nm). The retention factors (*R_f_*) of the enzymatic reaction products were calculated for riboflavin and FMN by

$$R_{f}= \frac{Distance traveled by component}{Distance traveled by solvent}$$

**Circular Dichroism spectroscopy of WT-*Mj*RibK and [*p*-HBI]-loop-*Mj*RibK.** The circular dichroism (CD) spectroscopy of WT-*Mj*RibK and [*p*-HBI]-loop-*Mj*RibK was conducted using a Jasco J-815 CD spectrophotometer. The measurements were performed in the far UV range (190 - 280 nm) using a quartz cuvette of path length 1 mm with a minimum of 3 accumulations. The data fitting and secondary structure content was estimated using the BeSTSel online tool.^4^

**UV-Visible Absorption and Emission Spectra of WT-*Mj*RibK and [*p*-HBI]-loop-*Mj*RibK.** Absorption and emission spectra were measured using a BioTek Cytation 5 multimode plate reader. Absorption spectra from 150 µL each of [*p*-HBI]-loop-*Mj*RibK (20 µM) and WT-*Mj*RibK (20 µM) were measured between 300 and 700 nm with a step size of 5 nm in a 96-well UV-transparent microplate. Emission spectra from 150 µL Red-*Mj*RibK (8 µM) at different excitation wavelengths (300, 325, 350, 400, 425, 450, 475, 500, and 525 nm) in a 96-well black polystyrene microplate with a fluorescence gain set at 100. The emission spectra were recorded up to 700 nm.

**Temperature Dependent Fluorescence Emission of [*p*-HBI]-loop-*Mj*RibK and Cyclability.** Emission spectra from 150 µL [*p*-HBI]-loop-*Mj*RibK (8 µM) at an excitation wavelength of 400 nm in a 96-well black polystyrene microplate with a fluorescence gain set at 100 was measured using a BioTek Cytation 5 microplate reader. The relative fluorescence units at 580 nm were measured at different temperatures (25, 35, 45, 55, and 65 °C). To assess the cyclability capability of [*p*-HBI]-loop-*Mj*RibK, the same sample underwent five cycles of heating and cooling, ranging from 25 °C to 65 °C. The corresponding relative fluorescence units (RFU) at 580 nm were recorded after reaching extreme temperature conditions (25 °C and 65 °C). To account for any volume changes resulting from evaporation at higher temperatures during the experiments, the sample volume was adjusted with Milli-Q water before each cycle.

**Fluorescence Lifetime of [*p*-HBI]-loop-*Mj*RibK**

The fluorescence lifetime in the time domain was determined using a home-built confocal microscope integrated with a lifetime measurement system. The system was equipped with LDH-D-C-485 nm pulsed LASER source (PicoQuant GmbH) for the fluorophore excitation. The fluorescence decay was detected by a Single-Photon Avalanche Diode (SPAD) solid-state detector (Excelitas Technologies®) connected to a TimeHarp 260 board (Time-Correlated Single Photon Counting (TCSPC) and Multi-Channel Scaling (MCS) board. Red-*Mj*RibK (10 µL) was added on a glass slide and placed on the sample stage of the confocal microscope. The protein sample was excited with a 485 nm pulse laser operating with a repetition rate of 8 MHz while the fluorescence decay was captured. Before the experiment, the laser power (100 µW) was measured at the objective. The acquired data was subsequently plotted and fitted using Origin® 9.0 software, employing the appropriate equation:

$$I_{t} = I_{0}e^{-(t/\tau)}$$

Where *I_o_* is the initial fluorescence intensity (time *t* = 0 µs), *I_t_* is the fluorescence intensity at time *t* µs, and *𝜏* is the fluorescence lifetime of the fluorophore.

**Molecular Dynamics Simulations**

**Force Field generation.** The initial conformation of *Mj*RibK was obtained from the protein data bank (PDB:2VBV). The water molecules associated with the protein were deleted from the PDB before further processing. The conformation of *p*-HBI (chromophore) in the negatively charged state, was generated from the PDB ID 7DNA. The restricted electrostatic potential (RESP) charges of *p*-HBI were obtained via the R.E.D. tool using the HF/6-31G* basis set.^5^^,^ ^6^ To simulate the protein environment surrounding *p*-HBI, the terminal of *p*-HBI was capped with N-methyl. The total charge of *p*-HBI was set to -1 and the charge on N-Methyl was set to 0 during charge calculations. Thereafter, ‘General Amber Force Field’ (GAFF) was used to generate the force field parameters of *p*-HBI using the calculated restrained electrostatic potential (RESP) charges.^7^ Finally, the initial conformation and the force field of the entire conjugated system including *p*-HBI, loop, and *Mj*RibK were generated using XLeap module of AmberTools, with the GAFF force field selected for *p*-HBI and the Amberff14sb force field for the loop-*Mj*RibK.^8-10^ The AMBER formatted force field and coordinates were converted to the GROMACS formatted coordinates and topology using a Python script ‘acpype.py’ (https://github.com/t-/acpype).^11^ All the simulation is performed using GROMACS formatted coordinates and topology using GROMACS-2020.2 version.^12^

**Simulation Details.** The [*p*-HBI]-loop-*Mj*RibK was placed in a cubic box of ~84 Å dimension. To eliminate any steric contact from the system an energy minimization in vacuum was performed using the steepest descent method for 10000 steps with the tolerance force 0.001 kJ mol^-1^.^13^ The system was solvated using the TIP3P water model, followed by the addition of 150 mM NaCl ions to neutralize the system.^14^^,^ ^15^ A second energy minimization was performed using the steepest descent method for 5,000 steps with a tolerance force of 0.001 kJ mol^-1^.^13^ Thereafter, the system was heated to 27 ºC using the Berendsen thermostat for 10 ns, during which the heavy atoms of the solute were positioned-restrained.^16^ To achieve a perfectly equilibrated system, annealing was performed for 100 ns. The annealing steps involved heating the system to 50 ºC in 3 steps and then cooling it back to 27 ºC in 3 steps. Subsequently, a 100 nanoseconds constant number of particles, temperature and pressure (NPT) simulation was performed at 27 ºC and 1 atm pressure using a Velocity-rescale thermostat and Berendsen barostat respectively, followed by a 1000 ns constant number of particles, volume and temperature (NVT) simulation at 300 K using Nose-Hoover thermostat.^16-19^ During the simulation, the bonds were constrained using the LINC algorithm.^20^ The electrostatic effects were treated with the Particle-mesh Ewald (PME) method using the 10 Å distance cut-off for long-range interaction.^21^

To get the statistically significant observation, two additional independent simulations were conducted using different initial structures (**Figure S8**) and velocities for a duration of 1 μs each. Subsequently, from all the trajectories, the final 500 ns simulation was extracted and merged to form a complete trajectory spanning a total length of 1.5 μs. All the analyses were performed on this comprehensive 1.5 μs trajectory. Furthermore, to examine the temperature dependence at 65 ºC on the system conformation, another 1 μs simulation was conducted using the starting conformation of **Figure S8 B**, while following the same protocol.

**Data Analysis.** All experiments were carried out independently three times unless otherwise mentioned. Data analyses were carried out using Microsoft Excel and Origin plotting suit. Data are represented as mean ± 1 standard deviation. All the MD simulation data was analyzed by using the various GROMACS modules and home-written code. The structural analyses of the proteins data bank (PDB) files were done using the visualization softwares 'UCSF Chimera' and 'Visual molecular dynamics (VMD).

**Table S1.** Nucleotide/amino acid sequence of the native proteins and fusion constructs. The sequence of KLGGGS linker is highlighted with red letters, and those of the fusion partners of PhoCl1 are highlighted in blue letters. The letters in violet are the C-terminal sequence of PhoCl1. The initial three amino acids H-Y-G are involved in the *p*-HBI chromophore formation of PhoCl1.

**PhoCl1-6xHis nucleotide sequence**

ATGGTGATCCCTGACTACTTCAAGCAGAGCTTCCCCGAGGGCTACAGCTGGGAGCGCAGCATGACCTACGAGGACGGCGGCATCTGCATCGCCACCAACGACATCACAATGGAGGGGGACAGCTTCATCAACAAGATCCACTTCAAGGGCACGAACTTCCCCCCCAACGGCCCCGTGATGCAGAAGAGGACCGTGGGCTGGGAGGCCAGCACCGAGAAGATGTACGAGCGCGACGGCGTGCTGAAGGGCGACGTGAAGATGAAGCTGCTGCTGAAGGGCGGCGGCCACTATCGCTGCGACTACCGCACCACCTACAAGGTCAAGCAGAAGCCCGTAAAGCTGCCCGACTACCACTTCGTGGACCACCGCATCGAGATCCTGAGCCACGACAAGGACTACAACAAGGTGAAGCTGTACGAGCACGCCGTGGCCCGCAACTCCACCGACAGCATGGACGAGCTGTACAAGGGTGGCAGCGGTGGCATGGTGAGCAAGGGCGAGGAGACCATTACAAGCGTGATCAAGCCTGACATGAAGAACAAGCTGCGCATGGAGGGCAACGTGAACGGCCACGCCTTCGTGATCGAGGGCGAGGGCAGCGGCAAGCCCTTCGAGGGCATCCAGACGATTGATTTGGAGGTGAAGGAGGGCGCCCCGCTGCCCTTCGCCTACGACATCCTGACCACCGCCTTCCACTACGGCAACCGCGTGTTCACCAAGTACCCACGGGGAGGTGGAGGTCTCGAGCACCACCACCACCACCACTGA

**PhoCl1-6xHis amino acid sequence**

MVIPDYFKQSFPEGYSWERSMTYEDGGICIATNDITMEGDSFINKIHFKGTNFPPNGPVMQKRTVGWEASTEKMYERDGVLKGDVKMKLLLKGGGHYRCDYRTTYKVKQKPVKLPDYHFVDHRIEILSHDKDYNKVKLYEHAVARNSTDSMDELYKGGSGGMVSKGEETITSVIKPDMKNKLRMEGNVNGHAFVIEGEGSGKPFEGIQTIDLEVKEGAPLPFAYDILTTAF**HYGNRVFTKYPR**GGGGLEHHHHHH

**WT-*Mj*RibK nucleotide sequence**

TTGGTGAAATTGATGATTATTGAGGGAGAAGTAGTTTCAGGACTTGGAGAAGGGAGATATTTTTTATCCCTCCCTCCTTACAAAGAGATATTTAAGAAGATTCTTGGCTTTGAACCTTATGAGGGGACATTAAATTTAAAATTAGATAGAGAATTTGATATAAACAAATTTAAATATATTGAAACAGAGGATTTTGAATTTAATGGGAAAAGATTTTTTGGAGTTAAGGTTTTACCAATAAAAATATTAATAGGTAATAAAAAAATAGATGGGGCGATAGTTGTGCCGAAAAAAACATATCATAGTAGTGAGATTATAGAGATAATTGCCCCAATGAAACTTAGGGAGCAATTTAATTTAAAGGATGGAGATGTTATAAAAATACTAATTAAGGGAGATAAAGATGAATAA

**WT-*Mj*RibK amino acid sequence**

MGSSHHHHHHSSGLVPRGSHMVKLMIIEGEVVSGLGEGRYFLSLPPYKEIFKKILGFEPYEGTLNLKLDREFDINKFKYIETEDFEFNGKRFFGVKVLPIKILIGNKKIDGAIVVPKKTYHSSEIIEIIAPMKLREQFNLKDGDVIKILIKGDKDE

**6xHis-(-30)GFP nucleotide sequence**

ATGGGTCATCACCACCACCATCACGGTGGCGCTAGCAAAGGTGAAGAGCTGTTTGACGGTGTAGTACCGATCTTAGTGGAATTAGACGGCGACGTGAACGGTCACGAATTTAGCGTGCGCGGCGAGGGCGAAGGTGACGCTACCGAGGGTGAATTGACCCTGAAGTTTATTTGCACAACAGGCGAATTACCCGTTCCGTGGCCCACCTTAGTGACCACCCTGACCTATGGCGTTCAGTGCTTCAGTGATTACCCAGATCATATGGATCAACACGATTTTTTCAAATCAGCCATGCCTGAAGGATATGTTCAAGAGCGTACAATCAGCTTCAAGGACGATGGCACCTATAAAACGCGTGCGGAAGTGAAATTTGAAGGCGACACATTAGTAAACCGTATCGAACTGAAAGGTATCGACTTCAAAGAAGACGGCAACATTTTAGGCCATAAGCTGGAATATAACTTTAATTCTCATGACGTGTATATTACGGCCGATAAACAGGAAAACGGTATCAAGGCAGAATTTGAAATTCGCCATAACGTGGAGGACGGCAGCGTTCAATTAGCGGATCATTATCAACAAAACACGCCGATTGGTGATGGGCCTGTACTGTTACCTGACGATCACTACCTGAGCACGGAGTCAGCCCTGAGCAAAGATCCGAACGAAGACCGCGATCACATGGTTCTGTTAGAATTCGTGACCGCTGCAGGCATTGATCATGGAATGGACGAGCTGTACAAGTAA

**6xHis-(-30)GFP amino acid sequence**

MGHHHHHHGGASKGEELFDGVVPILVELDGDVNGHEFSVRGEGEGDATEGELTLKFICTTGELPVPWPTLVTTLTYGVQCFSDYPDHMDQHDFFKSAMPEGYVQERTISFKDDGTYKTRAEVKFEGDTLVNRIELKGIDFKEDGNILGHKLEYNFNSHDVYITADKQENGIKAEFEIRHNVEDGSVQLADHYQQNTPIGDGPVLLPDDHYLSTESALSKDPNEDRDHMVLLEFVTAAGIDHGMDELYK

**6xHis-(Pos15)GFP nucleotide sequence**

ATGGGTCATCACCACCACCATCACGGTGGCGCTAGCAAAGGTGAACGTCTGTTTACGGGTGTAGTACCGATCTTAGTGGAATTAGACGGCGACGTGAACGGTCACAAATTTAGCGTGCGCGGCGAAGGCGAAGGTGACGCTACCCGTGGTAAATTGACCCTGAAGTTTATTTGCACAACAGGCAAATTACCCGTTCCGTGGCCCACCTTAGTGACCACCCTGACCTATGGCGTTCAGTGCTTCAGTCGTTACCCTAAACATATGAAACGTCACGATTTTT

TCAAATCAGCCATGCCTGAAGGATATGTTCAAGAGCGTACAATCAGCTTCAAGAAGGATGGCACCTATAAAACGCGTGCGGAAGTGAAATTTGAAGGCCGCACATTAGTAAACCGTATCGAACTGAAAGGTCGTGACTTCAAAGAAAAAGGCAACATTTTAGGCCATAAGCTGGAATATAACTTTAATTCTCATAACGTGTATATTACGGCCGATAAACGCAAGAATGGTATCAAGGCAAATTTCAAAATTCGCCATAACGTGAAAGACGGCAGCGTTCAATTAGCGGATCATTATCAACAAAACACGCCGATTGGTCGCGGGCCTGTACTGTTACCTCGCAACCACTACCTGAGCACCCGTTCAGCACTGAGCAAAGATCCGAAAGAAAAACGCGATCACATGGTTCTGTTAGAATTCGTGACCGCTGCAGGCATTACTCACGGAATGGACGAACTCTACAAGTAA

**6xHis-(Pos15)GFP amino acid sequence**

MGHHHHHHGGASKGERLFTGVVPILVELDGDVNGHKFSVRGEGEGDATRGKLTLKFICTTGKLPVPWPTLVTTLTYGVQCFSRYPKHMKRHDFFKSAMPEGYVQERTISFKKDGTYKTRAEVKFEGRTLVNRIELKGRDFKEKGNILGHKLEYNFNSHNVYITADKRKNGIKANFKIRHNVKDGSVQLADHYQQNTPIGRGPVLLPRNHYLSTRSALSKDPKEKRDHMVLLEFVTAAGITHGMDELYK

**6xHis-PhoCl1-*Mj*RibK fusion construct nucleotide sequence**

ATGGGCAGCAGCCATCATCATCATCATCACGGCGGTACCGTGATCCCTGACTACTTCAAGCAGAGCTTCCCCGAGGGCTACAGCTGGGAGCGCAGCATGACCTACGAGGACGGCGGCATCTGCATCGCCACCAACGACATCACAATGGAGGGGGACAGCTTCATCAACAAGATCCACTTCAAGGGCACGAACTTCCCCCCCAACGGCCCCGTGATGCAGAAGAGGACCGTGGGCTGGGAGGCCAGCACCGAGAAGATGTACGAGCGCGACGGCGTGCTGAAGGGCGACGTGAAGATGAAGCTGCTGCTGAAGGGCGGCGGCCACTATCGCTGCGACTACCGCACCACCTACAAGGTCAAGCAGAAGCCCGTAAAGCTGCCCGACTACCACTTCGTGGACCACCGCATCGAGATCCTGAGCCACGACAAGGACTACAACAAGGTGAAGCTGTACGAGCACGCCGTGGCCCGCAACTCCACCGACAGCATGGACGAGCTGTACAAGGGTGGCAGCGGTGGCATGGTGAGCAAGGGCGAGGAGACCATTACAAGCGTGATCAAGCCTGACATGAAGAACAAGCTGCGCATGGAGGGCAACGTGAACGGCCACGCCTTCGTGATCGAGGGCGAGGGCAGCGGCAAGCCCTTCGAGGGCATCCAGACGATTGATTTGGAGGTGAAGGAGGGCGCCCCGCTGCCCTTCGCCTACGACATCCTGACCACCGCCTTCCACTACGGCAACCGCGTGTTCACCAAGTACCCACGGAAGCTTGGCGGCGGCTCTTTGGTGAAATTGATGATTATTGAGGGAGAAGTAGTTTCAGGACTTGGAGAAGGGAGATATTTTTTATCCCTCCCTCCTTACAAAGAGATATTTAAGAAGATTCTTGGCTTTGAACCTTATGAGGGGACATTAAATTTAAAATTAGATAGAGAATTTGATATAAACAAATTTAAATATATTGAAACAGAGGATTTTGAATTTAATGGGAAAAGATTTTTTGGAGTTAAGGTTTTACCAATAAAAATATTAATAGGTAATAAAAAAATAGATGGGGCGATAGTTGTGCCGAAAAAAACATATCATAGTAGTGAGATTATAGAGATAATTGCCCCAATGAAACTTAGGGAGCAATTTAATTTAAAGGATGGAGATGTTATAAAAATACTAATTAAGGGAGATAAAGATGAATAA

**6xHis-PhoCl1-*Mj*RibK fusion amino acid sequence**

MGSSHHHHHHGGTVIPDYFKQSFPEGYSWERSMTYEDGGICIATNDITMEGDSFINKIHFKGTNFPPNGPVMQKRTVGWEASTEKMYERDGVLKGDVKMKLLLKGGGHYRCDYRTTYKVKQKPVKLPDYHFVDHRIEILSHDKDYNKVKLYEHAVARNSTDSMDELYKGGSGGMVSKGEETITSVIKPDMKNKLRMEGNVNGHAFVIEGEGSGKPFEGIQTIDLEVKEGAPLPFAYDILTTAF**HYGNRVFTKYPR**KLGGGSLVKLMIIEGEVVSGLGEGRYFLSLPPYKEIFKKILGFEPYEGTLNLKLDREFDINKFKYIETEDFEFNGKRFFGVKVLPIKILIGNKKIDGAIVVPKKTYHSSEIIEIIAPMKLREQFNLKDGDVIKILIKGDKDE

**6xHis-PhoCl1-(-30)GFP fusion nucleotide sequence**

ATGGGCAGCAGCCATCATCATCATCATCACGGCGGTACCGTGATCCCTGACTACTTCAAGCAGAGCTTCCCCGAGGGCTACAGCTGGGAGCGCAGCATGACCTACGAGGACGGCGGCATCTGCATCGCCACCAACGACATCACAATGGAGGGGGACAGCTTCATCAACAAGATCCACTTCAAGGGCACGAACTTCCCCCCCAACGGCCCCGTGATGCAGAAGAGGACCGTGGGCTGGGAGGCCAGCACCGAGAAGATGTACGAGCGCGACGGCGTGCTGAAGGGCGACGTGAAGATGAAGCTGCTGCTGAAGGGCGGCGGCCACTATCGCTGCGACTACCGCACCACCTACAAGGTCAAGCAGAAGCCCGTAAAGCTGCCCGACTACCACTTCGTGGACCACCGCATCGAGATCCTGAGCCACGACAAGGACTACAACAAGGTGAAGCTGTACGAGCACGCCGTGGCCCGCAACTCCACCGACAGCATGGACGAGCTGTACAAGGGTGGCAGCGGTGGCATGGTGAGCAAGGGCGAGGAGACCATTACAAGCGTGATCAAGCCTGACATGAAGAACAAGCTGCGCATGGAGGGCAACGTGAACGGCCACGCCTTCGTGATCGAGGGCGAGGGCAGCGGCAAGCCCTTCGAGGGCATCCAGACGATTGATTTGGAGGTGAAGGAGGGCGCCCCGCTGCCCTTCGCCTACGACATCCTGACCACCGCCTTCCACTACGGCAACCGCGTGTTCACCAAGTACCCACGGAAGCTTGGCGGCGGCTCTGGCGCTAGCAAAGGTGAAGAGCTGTTTGACGGTGTAGTACCGATCTTAGTGGAATTAGACGGCGACGTGAACGGTCACGAATTTAGCGTGCGCGGCGAGGGCGAAGGTGACGCTACCGAGGGTGAATTGACCCTGAAGTTTATTTGCACAACAGGCGAATTACCCGTTCCGTGGCCCACCTTAGTGACCACCCTGACCTATGGCGTTCAGTGCTTCAGTGATTACCCAGATCATATGGATCAACACGATTTTTTCAAATCAGCCATGCCTGAAGGATATGTTCAAGAGCGTACAATCAGCTTCAAGGACGATGGCACCTATAAAACGCGTGCGGAAGTGAAATTTGAAGGCGACACATTAGTAAACCGTATCGAACTGAAAGGTATCGACTTCAAAGAAGACGGCAACATTTTAGGCCATAAGCTGGAATATAACTTTAATTCTCATGACGTGTATATTACGGCCGATAAACAGGAAAACGGTATCAAGGCAGAATTTGAAATTCGCCATAACGTGGAGGACGGCAGCGTTCAATTAGCGGATCATTATCAACAAAACACGCCGATTGGTGATGGGCCTGTACTGTTACCTGACGATCACTACCTGAGCACGGAGTCAGCCCTGAGCAAAGATCCGAACGAAGACCGCGATCACATGGTTCTGTTAGAATTCGTGACCGCTGCAGGCATTGATCATGGAATGGACGAGCTGTACAAGTAA

**6xHis-PhoCl1-(-30)GFP fusion amino acid sequence**

MGSSHHHHHHGGTVIPDYFKQSFPEGYSWERSMTYEDGGICIATNDITMEGDSFINKIHFKGTNFPPNGPVMQKRTVGWEASTEKMYERDGVLKGDVKMKLLLKGGGHYRCDYRTTYKVKQKPVKLPDYHFVDHRIEILSHDKDYNKVKLYEHAVARNSTDSMDELYKGGSGGMVSKGEETITSVIKPDMKNKLRMEGNVNGHAFVIEGEGSGKPFEGIQTIDLEVKEGAPLPFAYDILTTAF**HYGNRVFTKYPR**KLGGGSGASKGEELFDGVVPILVELDGDVNGHEFSVRGEGEGDATEGELTLKFICTTGELPVPWPTLVTTLTYGVQCFSDYPDHMDQHDFFKSAMPEGYVQERTISFKDDGTYKTRAEVKFEGDTLVNRIELKGIDFKEDGNILGHKLEYNFNSHDVYITADKQENGIKAEFEIRHNVEDGSVQLADHYQQNTPIGDGPVLLPDDHYLSTESALSKDPNEDRDHMVLLEFVTAAGIDHGMDELYK

**6xHis-PhoCl1-(Pos15)GFP fusion nucleotide sequence**

AAGATCCACTTCAAGGGCACGAACTTCCCCCCCAACGGCCCCGTGATGCAGAAGAGGACCGTGGGCTGGGAGGCCAGCACCGAGAAGATGTACGAGCGCGACGGCGTGCTGAAGGGCGACGTGAAGATGAAGCTGCTGCTGAAGGGCGGCGGCCACTATCGCTGCGACTACCGCACCACCTACAAGGTCAAGCAGAAGCCCGTAAAGCTGCCCGACTACCACTTCGTGGACCACCGCATCGAGATCCTGAGCCACGACAAGGACTACAACAAGGTGAAGCTGTACGAGCACGCCGTGGCCCGCAACTCCACCGACAGCATGGACGAGCTGTACAAGGGTGGCAGCGGTGGCATGGTGAGCAAGGGCGAGGAGACCATTACAAGCGTGATCAAGCCTGACATGAAGAACAAGCTGCGCATGGAGGGCAACGTGAACGGCCACGCCTTCGTGATCGAGGGCGAGGGCAGCGGCAAGCCCTTCGAGGGCATCCAGACGATTGATTTGGAGGTGAAGGAGGGCGCCCCGCTGCCCTTCGCCTACGACATCCTGACCACCGCCTTCCACTACGGCAACCGCGTGTTCACCAAGTACCCACGGAAGCTTGGCGGCGGCTCTGGCGCTAGCAAAGGTGAACGTCTGTTTACGGGTGTAGTACCGATCTTAGTGGAATTAGACGGCGACGTGAACGGTCACAAATTTAGCGTGCGCGGCGAAGGCGAAGGTGACGCTACCCGTGGTAAATTGACCCTGAAGTTTATTTGCACAACAGGCAAATTACCCGTTCCGTGGCCCACCTTAGTGACCACCCTGACCTATGGCGTTCAGTGCTTCAGTCGTTACCCTAAACATATGAAACGTCACGATTTTTTCAAATCAGCCATGCCTGAAGGATATGTTCAAGAGCGTACAATCAGCTTCAAGAAGGATGGCACCTATAAAACGCGTGCGGAAGTGAAATTTGAAGGCCGCACATTAGTAAACCGTATCGAACTGAAAGGTCGTGACTTCAAAGAAAAAGGCAACATTTTAGGCCATAAGCTGGAATATAACTTTAATTCTCATAACGTGTATATTACGGCCGATAAACGCAAGAATGGTATCAAGGCAAATTTCAAAATTCGCCATAACGTGAAAGACGGCAGCGTTCAATTAGCGGATCATTATCAACAAAACACGCCGATTGGTCGCGGGCCTGTACTGTTACCTCGCAACCACTACCTGAGCACCCGTTCAGCACTGAGCAAAGATCCGAAAGAAAAACGCGATCACATGGTTCTGTTAGAATTCGTGACCGCTGCAGGCATTACTCACGGAATGGACGAACTCTACAAGTAA

**6xHis-PhoCl1-(Pos15)GFP fusion amino acid sequence**

MGSSHHHHHHGGTVIPDYFKQSFPEGYSWERSMTYEDGGICIATNDITMEGDSFINKIHFKGTNFPPNGPVMQKRTVGWEASTEKMYERDGVLKGDVKMKLLLKGGGHYRCDYRTTYKVKQKPVKLPDYHFVDHRIEILSHDKDYNKVKLYEHAVARNSTDSMDELYKGGSGGMVSKGEETITSVIKPDMKNKLRMEGNVNGHAFVIEGEGSGKPFEGIQTIDLEVKEGAPLPFAYDILTTAF**HYGNRVFTKYPR**KLGGGSGASKGERLFTGVVPILVELDGDVNGHKFSVRGEGEGDATRGKLTLKFICTTGKLPVPWPTLVTTLTYGVQCFSRYPKHMKRHDFFKSAMPEGYVQERTISFKKDGTYKTRAEVKFEGRTLVNRIELKGRDFKEKGNILGHKLEYNFNSHNVYITADKRKNGIKANFKIRHNVKDGSVQLADHYQQNTPIGRGPVLLPRNHYLSTRSALSKDPKEKRDHMVLLEFVTAAGITHGMDELYK

**Table S2.** Primers used to amplify the gene products of PhoCl1, *Mj*RibK, and supercharged proteins. The forward and reverse primers have been indicated by the suffixes F and R respectively.

| **#** | **Primers** | **5’-3’ sequence** |
| --- | --- | --- |
| 1 | PhoCl1_F | GTTTAACTTTAAGAAGGAGATATACCATGGGCAGCAGCCATCATCATCATCATCACGGCGGTACCGTGATCCCTGACTACTTCAAGC |
| 2 | PhoCl1_R | AGAGCCGCCGCCAAGCTTCCGTGGGTACTTGGTGAAC |
| 3 | MjRibK_F | CGGAAGCTTGGCGGCGGCTCTTTGGTGAAATTGATGATTATTG |
| 4 | MjRibK_R | TCAGTGGTGGTGGTGGTGGTGCTCGAGTTATTCATCTTTATCTCCCTTAAT |
| 5 | (-30)GFP_F | GAAGCTTGGCGGCGGCTCTGGCGCTAGCAAAGGTGAAGAG |
| 6 | (-30)GFP_R^#^ | GTGGTGGTGGTGCTCGAGTTACTTGTACAGCTCGTCCATTCC |
| 7 | (Pos15)GFP_F | GAAGCTTGGCGGCGGCTCTGGCGCTAGCAAAGGTGAAC |
| 8 | (Pos15)GFP_R^#^ | GTGGTGGTGGTGCTCGAGTTACTTGTACAGCTCGTCCATTCC |

\# The reverse primers for the amplification of (-30)GFP, and (Pos15)GFP are same.

**Table S3.** Determination of the secondary structure content of [*p*-HBI]-loop-*Mj*RibK and WT-*Mj*RibK through circular dichroism spectroscopy.

| **Secondary structure** | **[*p*-HBI]-loop-*Mj*RibK (%)** | **WT-*Mj*RibK (%)** |
| --- | --- | --- |
| Helix | 15.7 | 18.7 |
| Antiparallel | 49.3 | 40.4 |
| Parallel | 19.2 | 25.3 |
| Turn | 15.8 | 15.6 |
| Others | 0 | 0 |
| Total | 100 | 100 |

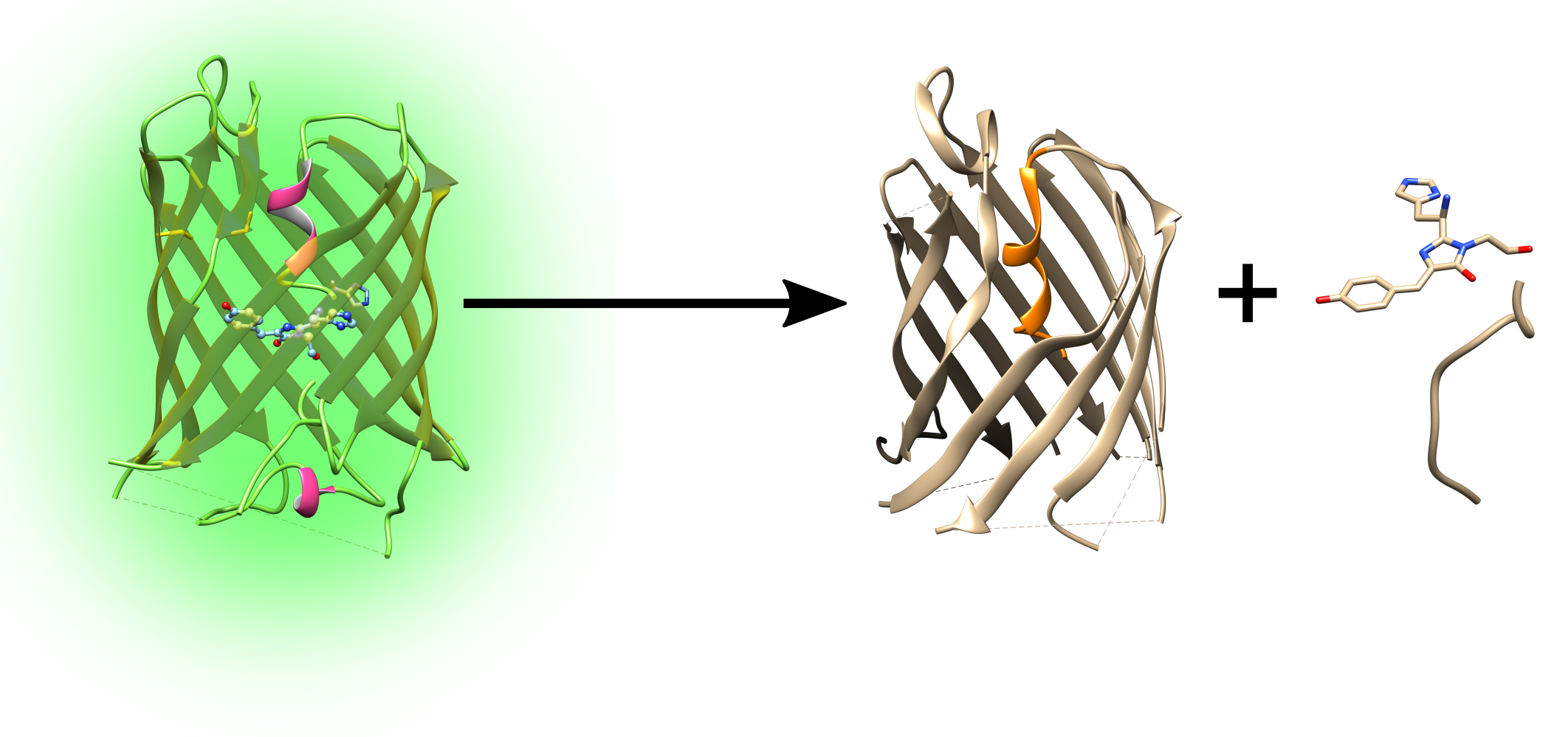

Photocleavable protein PhoCl1

Empty PhoCl1 barrel

C-terminal

fragment

([*p*-HBI]-NRVFTKYPR)

400 nm light

Photocleavage

**Figure S1.** A schematic of the photocleavage mechanism of the photocleavable protein (PhoCl1). The exposure of PhoCl1 to 400 nm light results in the fragmentation of PhoCl1 into two components. The first one is the PhoCl1 empty β-barrel and second is the C-terminal fragment containing the *p*-HBI chromophore followed by NRVFTKYPR amino acid sequence. Both fragments show non-fluorescent characteristics post photocleavage.

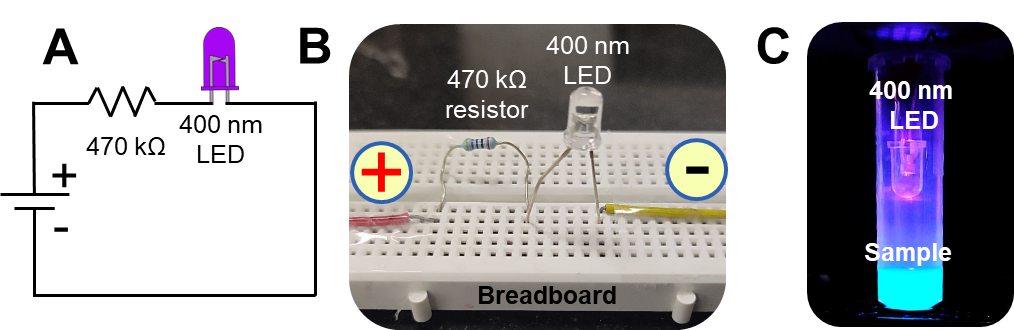

**Figure S2.** The custom designed 400 nm LED setup for the light induced cleavage of PhoCl, and its fusion protein constructs. **(A)** The circuit diagram of the custom built 400 nm LED setup. A 470 kΩ resistor was attached to the positive terminal of LED. The positive (red wire) and the negative (yellow wire) terminals were connected to the power source. **(B)** The image of a LED setup built on a breadboard. **(C)** The 400 nm LED setup in an inverted position over PhoCl1 (green liquid) kept inside the 2 mL microcentrifuge tube.

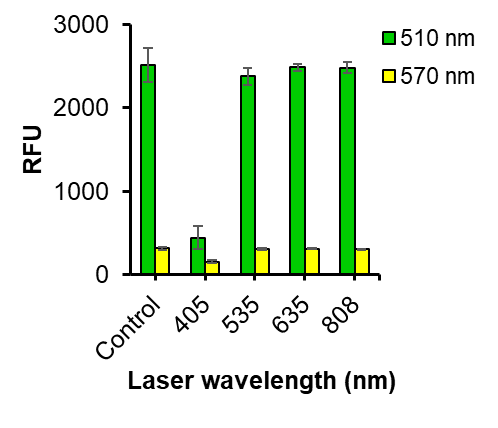

**Figure S3.** The relative fluorescence intensity of PhoCl1 after photoexposure to the LASERs of wavelengths 405, 535, 635 and 808 nm (~13 mWcm^-2^ for 405 nm LASER, ~65 mWcm^-2^ for others; Time = 1 h). The green bars indicate relative fluorescence intensities measured at 510 nm (green fluorescence). The yellow bars indicate relative fluorescence intensities measured at 570 nm (red fluorescence). The relative fluorescence intensity of the LASER exposed PhoCl1 was compared to the relative fluorescence intensity of the unexposed sample. A statistically significant decrease in the relative fluorescence intensity at 510 nm is observed when PhoCl1 was exposed to 405 nm while relative fluorescence intensity remains unchanged when exposed to other LASER wavelengths. No change in the relative fluorescence intensity at 570 nm is observed when PhoCl1 was exposed to all wavelengths. These indicate PhoCl1 does not possess the ability to generate red fluorescence independently after photo exposure.

**400 nm LED**

(+)

(+)

(-)

(-)

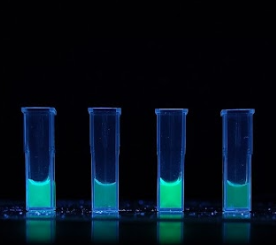

PhoCl1-GFP(-30)

PhoCl1-GFP(-30)

PhoCl1-GFP(+15)

PhoCl1-GFP(+15)

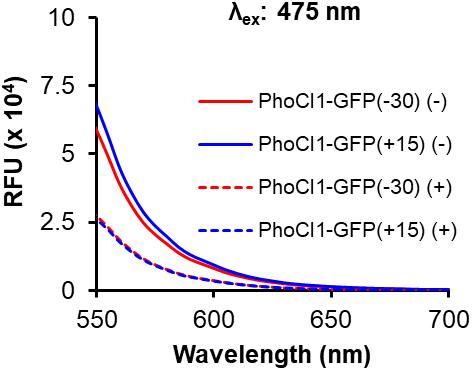

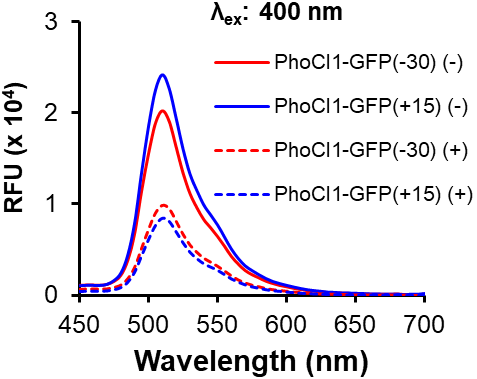

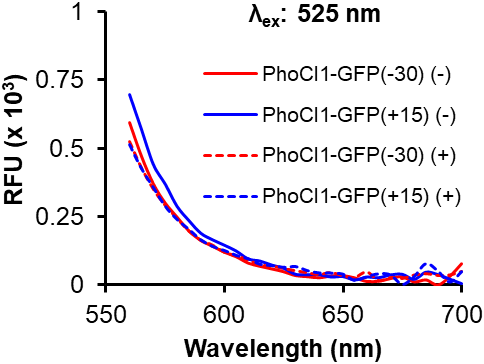

**B**

**A**

**C**

**D**

**E**

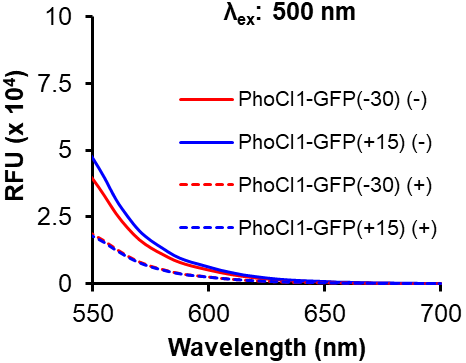

λex: 475 nm

RFU (x 104)

| Wavelength (nm) | PhoCl1-GFP(-30) (-) | PhoCl1-GFP(+15) (-) | PhoCl1-GFP(-30) (+) | PhoCl1-GFP(+15) (+) |
| --- | --- | --- | --- | --- |
| 540 | 76180.3 | 87906.7 | 34695.7 | 33171.3 |
| 545 | 67715.7 | 78075 | 30996.7 | 29617.7 |
| 550 | 59184.3 | 68084 | 27457 | 26183.7 |
| 555 | 49290.7 | 56941.3 | 23273 | 22305.7 |
| 560 | 38978.7 | 44948.7 | 18593.3 | 17800.7 |
| 565 | 31113.3 | 35966.3 | 14866.3 | 14219 |
| 570 | 24890.7 | 28823.3 | 11798.7 | 11277 |
| 575 | 20531 | 23742 | 9591 | 9238.33 |
| 580 | 17069.7 | 19793.7 | 7879 | 7552.33 |
| 585 | 13775.3 | 15909.3 | 6380 | 6056.33 |
| 590 | 11331.7 | 13069 | 5235 | 5008.67 |
| 595 | 9654.67 | 11133 | 4464.33 | 4279.67 |
| 600 | 8083.67 | 9348.67 | 3777.33 | 3592.67 |
| 605 | 6502.33 | 7566 | 3093.33 | 2944 |
| 610 | 5328.33 | 6171.33 | 2514 | 2419 |
| 615 | 4343.67 | 4993 | 2038 | 1976.33 |
| 620 | 3543.33 | 4094.67 | 1703.33 | 1600.33 |
| 625 | 2918 | 3422 | 1380.33 | 1334.33 |
| 630 | 2389.33 | 2767.33 | 1131.67 | 1068.33 |
| 635 | 1949.67 | 2275.67 | 915 | 886.667 |
| 640 | 1668 | 1940 | 794.667 | 777.333 |
| 645 | 1447.67 | 1649 | 663.667 | 659.667 |
| 650 | 1217.67 | 1377 | 560 | 539 |
| 655 | 993.667 | 1155 | 452 | 473.667 |
| 660 | 814 | 950.667 | 384 | 374 |
| 665 | 701.333 | 824.333 | 347.667 | 316.667 |
| 670 | 595.333 | 681 | 304.333 | 262.333 |
| 675 | 496.333 | 579 | 256.333 | 216.667 |
| 680 | 424 | 465 | 220.333 | 210.667 |
| 685 | 315 | 400 | 183 | 219.333 |
| 690 | 337.333 | 314 | 229.333 | 238 |
| 695 | 247.667 | 335.667 | 136 | 118 |
| 700 | 236 | 255.333 | 166.333 | 118.667 |

λex: 400 nm

RFU (x 104)

| Wavelength (nm) | PhoCl1-GFP(-30) (-) | PhoCl1-GFP(+15) (-) | PhoCl1-GFP(-30) (+) | PhoCl1-GFP(+15) (+) |
| --- | --- | --- | --- | --- |
| 450 | 1067 | 1032.33 | 686.333 | 454 |
| 455 | 1097.67 | 1071 | 724.333 | 482.667 |
| 460 | 1095.33 | 1069.33 | 720 | 478 |
| 465 | 1077 | 1063.67 | 708.333 | 472.667 |
| 470 | 1172.67 | 1188 | 739.667 | 508.667 |
| 475 | 1509.33 | 1610.67 | 891.333 | 641.667 |
| 480 | 2190.67 | 2434 | 1170.33 | 891.667 |
| 485 | 3846 | 4416.67 | 1861.67 | 1466 |
| 490 | 6848.67 | 8014.67 | 3134.67 | 2589.67 |
| 495 | 11465.3 | 13505.7 | 5197.67 | 4356.33 |
| 500 | 15525.3 | 18523 | 7245.67 | 6118.33 |
| 505 | 18864 | 22474.3 | 9027.67 | 7647.67 |
| 510 | 20209 | 24145 | 9871.67 | 8431.33 |
| 515 | 19023.3 | 22588.7 | 9486 | 8074.67 |
| 520 | 16216.3 | 19358.7 | 8120.33 | 6948.33 |
| 525 | 13371.3 | 15987.7 | 6696.33 | 5756.67 |
| 530 | 11133 | 13262.7 | 5529 | 4733.67 |
| 535 | 9577.67 | 11461.7 | 4739 | 4027.67 |
| 540 | 8279 | 9880.33 | 4086.33 | 3473.67 |
| 545 | 7372.33 | 8777.67 | 3659.33 | 3111.33 |
| 550 | 6419 | 7682 | 3214 | 2756 |
| 555 | 5351.33 | 6365 | 2694.33 | 2294.33 |
| 560 | 4235 | 5044 | 2162.33 | 1824.33 |
| 565 | 3392.33 | 4004 | 1711 | 1464 |
| 570 | 2696 | 3204.67 | 1374.33 | 1177.67 |
| 575 | 2235.33 | 2681.67 | 1144 | 956.667 |
| 580 | 1877 | 2226 | 961.333 | 799.333 |
| 585 | 1520 | 1816 | 774.333 | 653.667 |
| 590 | 1270.33 | 1487.33 | 648.333 | 533 |
| 595 | 1058 | 1257.33 | 559.333 | 472 |
| 600 | 907.667 | 1068.67 | 480.333 | 391.667 |
| 605 | 748.333 | 889 | 394.333 | 325.667 |
| 610 | 613.667 | 726.667 | 320.667 | 263.333 |
| 615 | 495.667 | 590 | 270 | 219.333 |
| 620 | 415 | 494 | 226.667 | 185.667 |
| 625 | 355 | 413.667 | 181.667 | 151.667 |
| 630 | 278.667 | 326 | 158.667 | 127 |
| 635 | 248 | 296.333 | 136.333 | 104.667 |
| 640 | 207.667 | 240 | 117 | 89.3333 |
| 645 | 184.333 | 227.667 | 108.333 | 73.6667 |
| 650 | 152.333 | 196.333 | 76.6667 | 65.3333 |
| 655 | 117.667 | 169.333 | 70 | 52.6667 |
| 660 | 92.3333 | 136.667 | 76 | 40 |
| 665 | 73.3333 | 126.333 | 42 | 64.6667 |
| 670 | 73.6667 | 95.3333 | 88.6667 | 28.3333 |
| 675 | 88 | 90.3333 | 81 | 39 |
| 680 | 115 | 111 | 65.6667 | 88.6667 |
| 685 | 67 | 51.3333 | 29.6667 | 11.6667 |
| 690 | 45.3333 | 103 | 35 | 23.6667 |
| 695 | 51.6667 | 87 | 0 | 27 |
| 700 | 87.3333 | 174 | 49.6667 | 50.6667 |

λex: 525 nm

RFU (x 103)

| Wavelength (nm) | PhoCl1-GFP(-30) (-) | PhoCl1-GFP(+15) (-) | PhoCl1-GFP(-30) (+) | PhoCl1-GFP(+15) (+) |
| --- | --- | --- | --- | --- |
| 560 | 594 | 697 | 524.333 | 515.333 |
| 565 | 471.333 | 569.667 | 428.667 | 428 |
| 570 | 370.333 | 440.667 | 350.667 | 358 |
| 575 | 300 | 368.667 | 291.333 | 289.667 |
| 580 | 248.667 | 287 | 243.667 | 240.667 |
| 585 | 198 | 234.667 | 197 | 204.667 |
| 590 | 163 | 188.333 | 163.667 | 166 |
| 595 | 142 | 164 | 145.333 | 141.667 |
| 600 | 120 | 144 | 119.667 | 126.667 |
| 605 | 103.333 | 122.667 | 106.333 | 108.667 |
| 610 | 80.3333 | 93.6667 | 91.3333 | 96.3333 |
| 615 | 68.6667 | 86 | 73 | 76.3333 |
| 620 | 59 | 71 | 69 | 71.3333 |
| 625 | 48.6667 | 59.3333 | 60 | 64.6667 |
| 630 | 34.6667 | 37.3333 | 51.6667 | 67.6667 |
| 635 | 30.6667 | 36 | 44.3333 | 52.6667 |
| 640 | 26.6667 | 36 | 38 | 44.3333 |
| 645 | 33.3333 | 27.3333 | 45.3333 | 43.6667 |
| 650 | 34.6667 | 29 | 32.6667 | 39.6667 |
| 655 | 26 | 16.6667 | 27.3333 | 17 |
| 660 | 13.3333 | 27 | 47.6667 | 30.6667 |
| 665 | 14 | 25 | 15.3333 | 41 |
| 670 | 23.6667 | 35 | 26.6667 | 37.3333 |
| 675 | 30.6667 | 37.3333 | 23.6667 | 2.33333 |
| 680 | 11.3333 | 19 | 36 | 31.3333 |
| 685 | 18.6667 | 45 | 41 | 78.3333 |
| 690 | 0 | 36.3333 | 34.6667 | 51.3333 |
| 695 | 31.6667 | 19.3333 | 36.3333 | 11 |
| 700 | 78 | 3 | 49.3333 | 49 |

λex: 500 nm

RFU (x 104)

| Wavelength (nm) | PhoCl1-GFP(-30) (-) | PhoCl1-GFP(+15) (-) | PhoCl1-GFP(-30) (+) | PhoCl1-GFP(+15) (+) |
| --- | --- | --- | --- | --- |
| 530 | 69592.3 | 83261.7 | 34784.3 | 33650 |
| 535 | 57800.3 | 69741.7 | 27331.3 | 26315 |
| 540 | 49608.3 | 59769 | 22954 | 22048.7 |
| 545 | 44197 | 53366.3 | 20492 | 19771 |
| 550 | 39132.3 | 47039 | 18467 | 17710 |
| 555 | 32880.7 | 39714 | 15947.3 | 15371 |
| 560 | 26278 | 31574.7 | 12978.7 | 12467.7 |
| 565 | 20923.3 | 25273.3 | 10365.7 | 9969.67 |
| 570 | 16571 | 19949.3 | 8188.67 | 7865.33 |
| 575 | 13503.7 | 16285.7 | 6560 | 6308 |
| 580 | 11122.3 | 13426.7 | 5363.33 | 5157 |
| 585 | 8950.33 | 10722.7 | 4280.33 | 4114.67 |
| 590 | 7389.67 | 8872 | 3487 | 3363 |
| 595 | 6265.67 | 7585.67 | 3003 | 2904.33 |
| 600 | 5251.33 | 6372 | 2564.33 | 2471 |
| 605 | 4329 | 5187 | 2078.67 | 2046 |
| 610 | 3533.67 | 4260.67 | 1731 | 1668.67 |
| 615 | 2871.33 | 3417.67 | 1388 | 1371.67 |
| 620 | 2344 | 2823.33 | 1139.67 | 1119 |
| 625 | 1927.67 | 2296.33 | 951.667 | 919.333 |
| 630 | 1566 | 1842.33 | 760 | 736.667 |
| 635 | 1263.33 | 1532.67 | 617.667 | 574.333 |
| 640 | 1078.33 | 1303.33 | 515 | 525.667 |
| 645 | 924.333 | 1108.33 | 455.667 | 436.667 |
| 650 | 767 | 944 | 379 | 377 |
| 655 | 644.667 | 751 | 308.333 | 309 |
| 660 | 520.333 | 672.667 | 284.333 | 238.333 |
| 665 | 443 | 560 | 222 | 241.667 |
| 670 | 376.333 | 448.667 | 190.333 | 185 |
| 675 | 327.667 | 366 | 126.667 | 136.333 |
| 680 | 273.333 | 361.667 | 159 | 124.333 |
| 685 | 203.667 | 302 | 123.333 | 135.667 |
| 690 | 227 | 284.667 | 94.3333 | 104.667 |
| 695 | 187.333 | 245 | 167 | 58 |
| 700 | 194.667 | 169.667 | 138 | 102 |

**Figure S4.** Photoexposure of fusion proteins of PhoCl1 with supercharged variants of green fluorescent protein to 400 nm light for 12 h **(A)** Digital images of PhoCl1-(-30)GFP and PhoCl1-(+15)GFP before and after photo exposure acquired under a UV-transilluminator. The (-) and (+) signs indicate unexposed and photoexposed sample respectively. A clear decrease in green fluorescence of the supercharged fusion variants is observed after photo exposure. Additionally, no evident red fluorescence is observable in both the unexposed and photo-exposed samples. **(B-E)** The comparison of emission spectra at different excitation wavelengths (400, 475, 500, and 525 nm) of the unexposed and photoexposed fusion proteins of PhoCl1 with supercharged variants of green fluorescent protein. A clear decrease in green fluorescence in all fusion constructs is observed after photoexposure. Additionally, no new peak corresponding to red fluorescence emerged indicating the requirement of *Mj*RibK as the fusion partner.

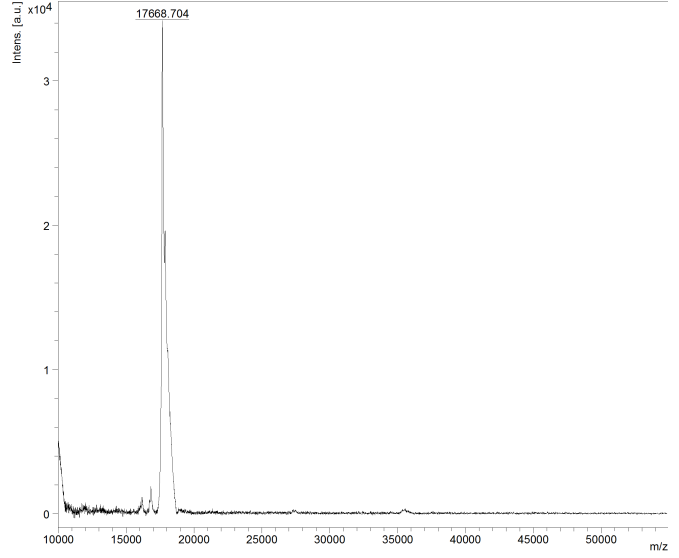

**Figure S5.** The MALDI-TOF mass spectrum of the [*p*-HBI]-loop-*Mj*RibK. The analysis displays a peak at m/z 17668.704 corresponding to the molecular weight of [*p*-HBI]-loop-*Mj*RibK (Theoretical mass: 17686.79 Da)

***Mj*RibK**

NS

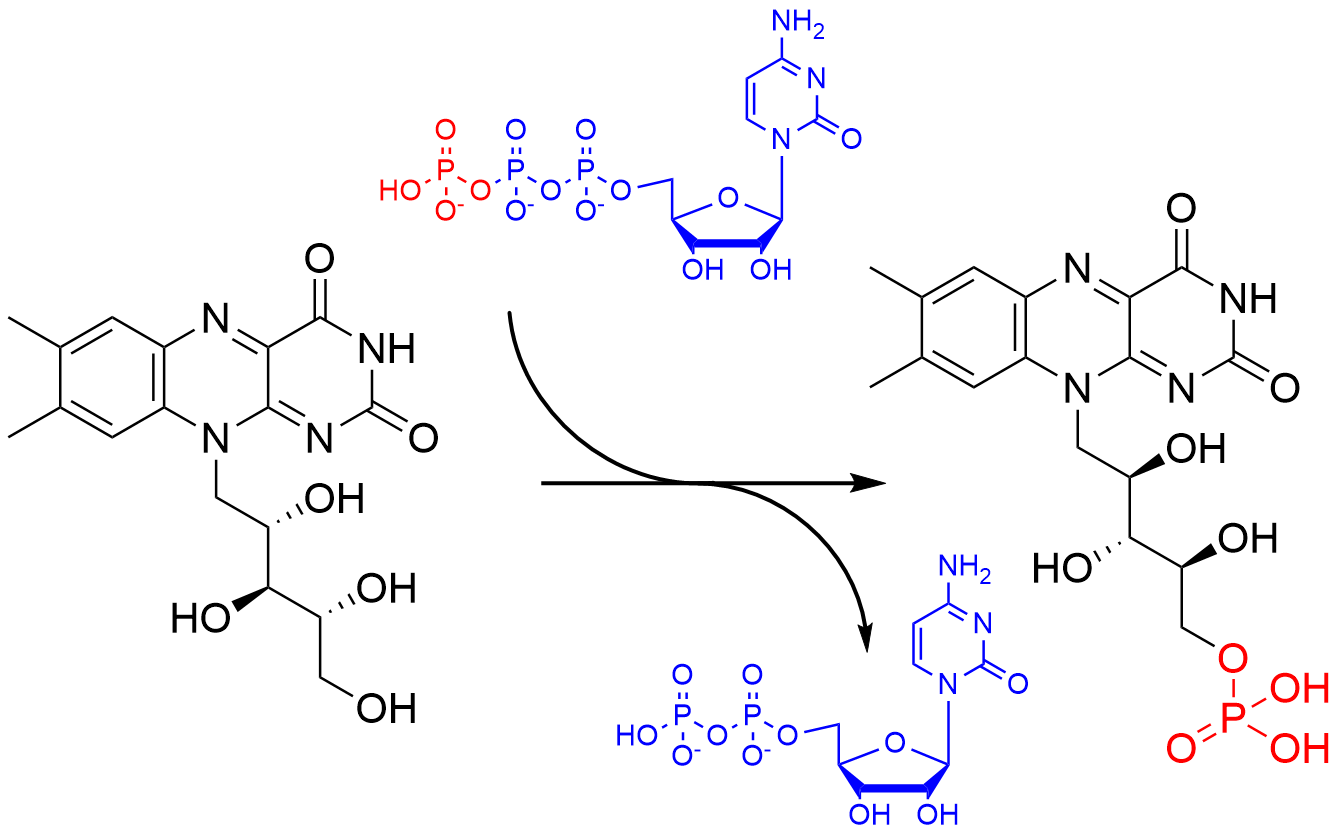

Riboflavin

FMN

CTP

CDP

**B**

**A**

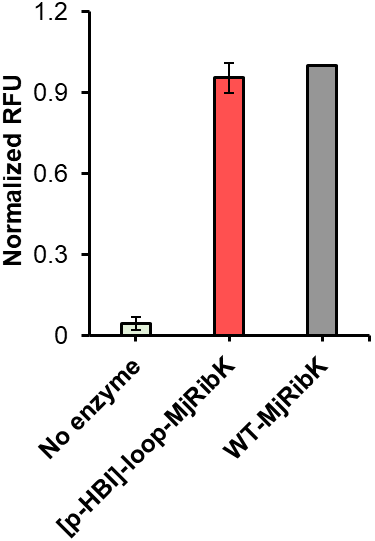

Normalized RFU

|  | Average |
| --- | --- |
| No enzyme | 0.0466045 |
| [p-HBI]-loop-MjRibK | 0.954585 |
| WT-MjRibK | 1 |

**Figure S6.** The enzyme activity of [*p*-HBI]-loop-*Mj*RibK and WT-*Mj*RibK through riboflavin kinase assay. **(A)** Reaction schematic of enzymatic conversion of riboflavin into flavin monophosphate (FMN) in the presence of *Mj*RibK using CTP as a phosphate donor **(B)** The riboflavin kinase assay was set up individually with WT-*Mj*RibK, [*p*-HBI]-loop-*Mj*RibK and ‘no enzyme control’. The reaction mixtures (1 µL) were spotted on a TLC plate. The mobile phase used to run the TLC was the upper layer of the mixture of solvent- 2-butanol, acetic acid, and water in 4:1:5 ratio. The TLC plate was dried and the fluorescent spots corresponding to the reactant ‘riboflavin’ and the product ‘FMN’ were captured on a UV transilluminator using long UV. The fluorescent spots were analyzed by densitometry using ImageJ. A two-tailed unpaired t-test between the activity of [*p*-HBI]-loop-*Mj*RibK and WT-*Mj*RibK displays no significant differences in activity (*p* = 0.23). These indicate minimal structural perturbations between [*p*-HBI]-loop-*Mj*RibK and WT-*Mj*RibK.

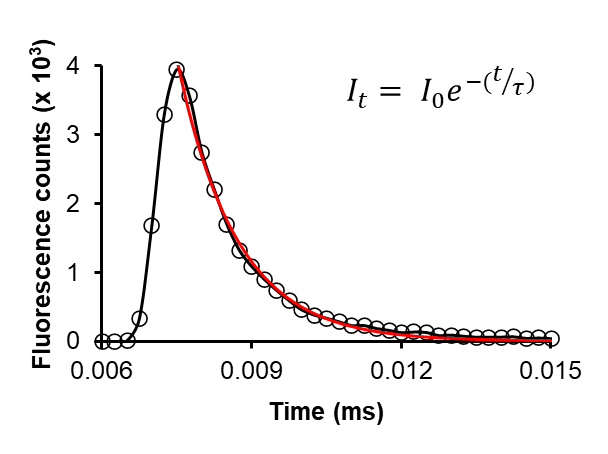

**Figure S7.** Fluorescence lifetime measurement of [*p*-HBI]-loop-*Mj*RibK. The exponential decay curve (black circles connected with a black solid line) was plotted and fitted (red solid line) with the fluorescence decay equation mentioned as an inset. The lifetime of [*p*-HBI]-loop-*Mj*RibK was determined to be 1.2 ns.

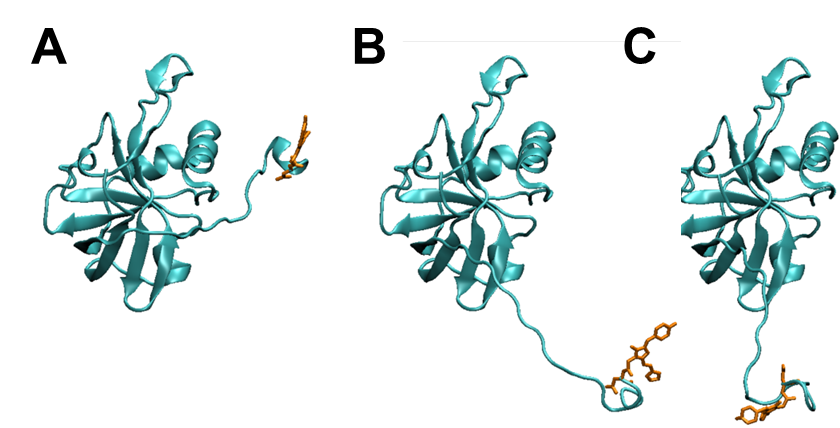

**Figure S8.** Initial conformations of *p*-HBI-loop-*Mj*RibK used for the three independent molecular dynamic simulations using different velocities. **(A)** Initial conformation 1 where the center of mass distance between the *Mj*RibK to *p-*HBI is 28.7 Å (B) Initial conformation 2 where the center of mass distance between the *Mj*RibK and *p-*HBI is 51.2 Å and (C) Initial conformation 3 where the center of mass distance between the *Mj*RibK to *p-*HBI is 53 Å. The protein is shown with a ribbon representation (dark cyan) and the *p*-HBI (light brown) is shown in a licorice representation.

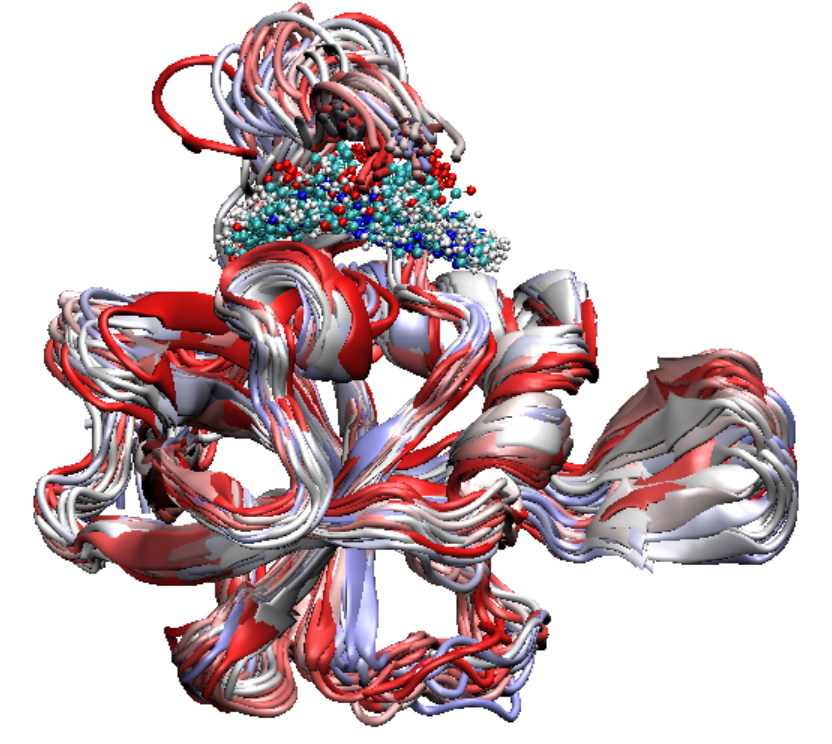

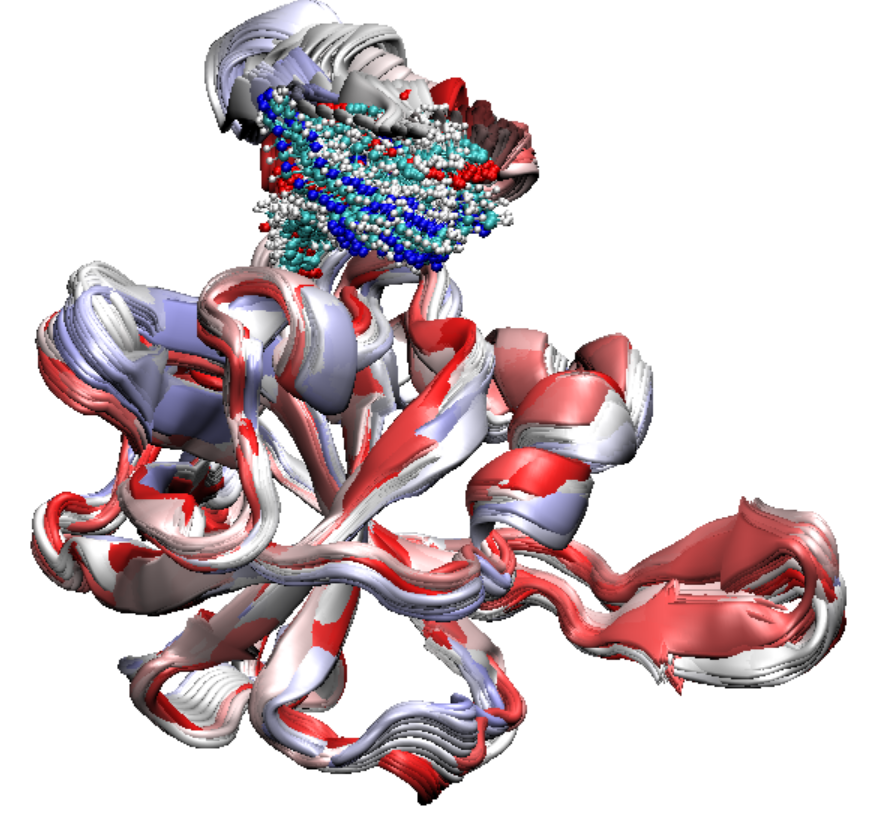

**A**

**B**

**Figure S9.** A cluster of 50 superimposed structures of [*p*-HBI]-loop-*Mj*RibK in (A) C2 state and (B) C3 state**.** The 50 representative structures are superimposed to illustrate the differences in conformation between the C2 and C3 states. The protein is represented as a ribbon in light green color, while *p*-HBI is represented as licorice. The greater dispersion in the position of *p-*HBI indicates increased fluctuation at the C3 state compared to the C2 state.

**Figure S10.** Total interaction energy (electrostatic and van der Waals) of individual amino acids of [*p*-HBI]-loop-*Mj*RibK protein with the chromophore *p*-HBI. The average interaction energy is calculated over 5000 frames of C1 state using the GROMACS energy module. All the residues with interaction energy lesser than -10 kJ mol^-1^ are listed in **Figure 3C**.

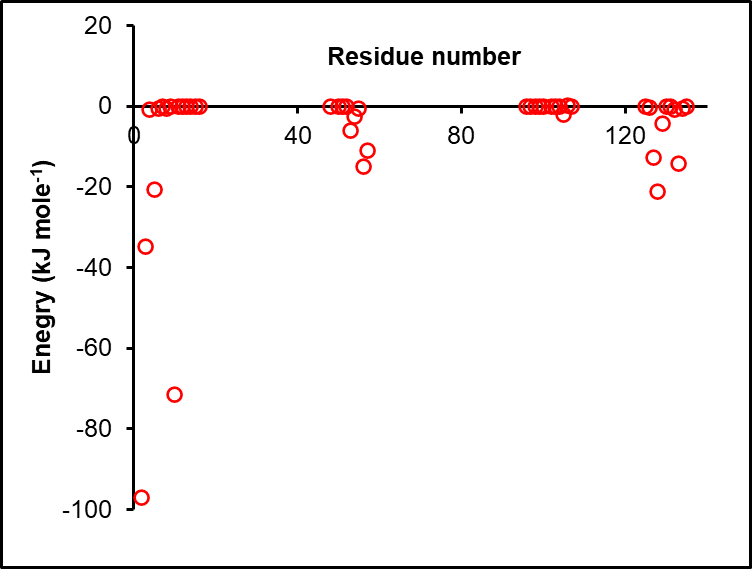

Asn2

Arg3

Arg10

Phe5

Tyr56

Glu57

Pro127

Gln133

Met128

| Residue number | Enegry (kJ mole-1) |
| --- | --- |
| 2 | -97.0719 |
| 3 | -34.7887 |
| 4 | -0.92375 |
| 5 | -20.6779 |
| 6 | -0.59377 |
| 7 | -0.02316 |
| 8 | -0.66549 |
| 9 | -0.01604 |
| 10 | -71.6179 |
| 11 | -0.00179 |
| 12 | -0.000452467 |
| 13 | -0.00255 |
| 14 | -0.00268 |
| 15 | -0.00205 |
| 16 | -5.9315e-05 |
| 48 | -0.00494 |
| 50 | -3.0185e-05 |
| 51 | -0.0622 |
| 52 | -0.03564 |
| 53 | -6.02784 |
| 54 | -2.50328 |
| 55 | -0.58938 |
| 56 | -14.9136 |
| 57 | -10.982 |
| 96 | 8.01983e-06 |
| 97 | -0.06379 |
| 98 | -0.00445 |
| 99 | -1.53943e-05 |
| 100 | -0.00343 |
| 102 | -0.00022806 |
| 103 | -0.11639 |
| 104 | -0.07242 |
| 105 | -1.99792 |
| 106 | 0.24181 |
| 107 | -0.00522 |
| 125 | -0.00179 |
| 126 | -0.39024 |
| 127 | -12.8662 |
| 128 | -21.2563 |
| 129 | -4.275 |
| 130 | -0.17395 |
| 131 | -0.00714 |
| 132 | -0.84768 |
| 133 | -14.1286 |
| 134 | -0.58857 |
| 135 | -0.00361 |

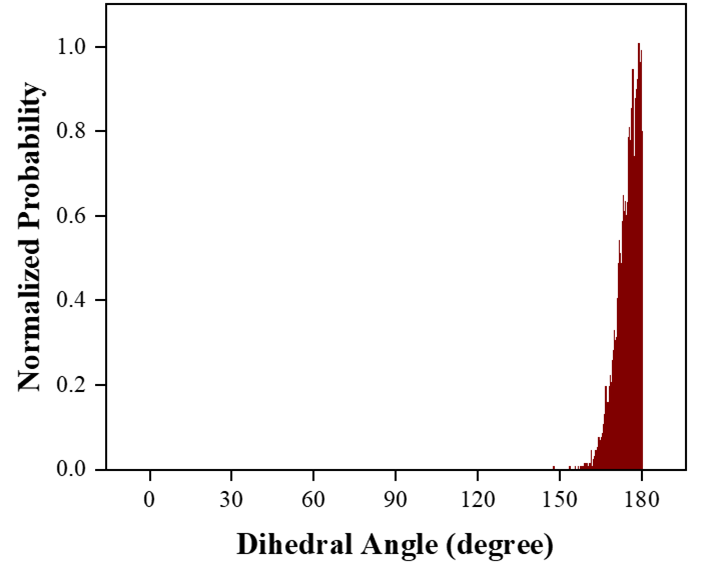

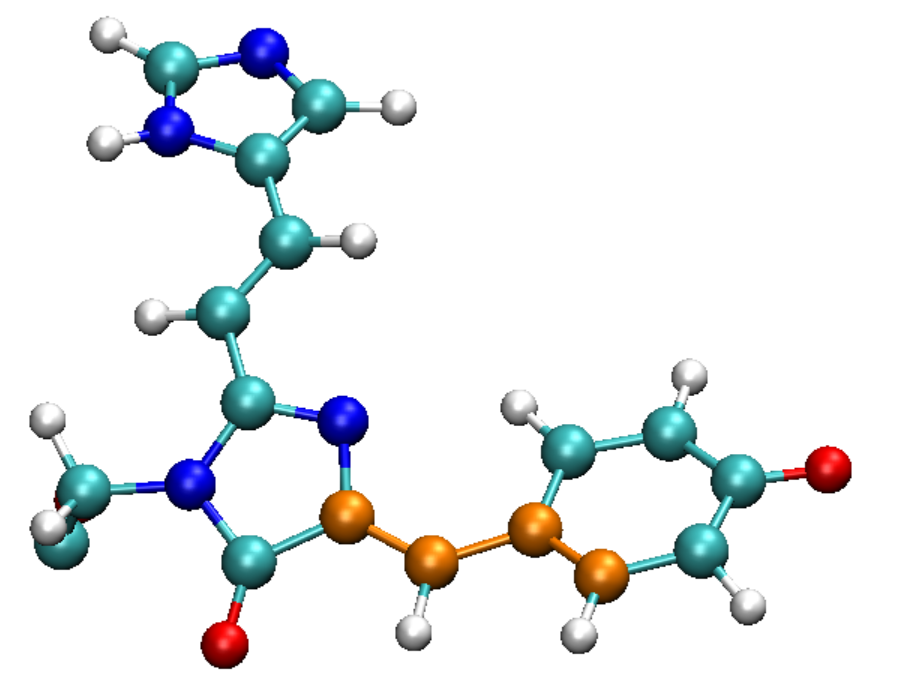

C1

C2

C3

C4

**Figure S11.** The plot shows the dihedral between the C1-C2-C3-C4 atoms in *p*-HBI. This revealed a population predominantly ranging between 171°-179° suggesting a nearly planar orientation of *p*-HBI in [*p*-HBI]-loop-*Mj*RibK.

|  | No. of water molecules around p-HBI |
| --- | --- |
| Unbound p-HBI (25°C) | 50 |
| Bound p-HBI (25 °C) | 12 |
| Bound p-HBI (65 °C) | 34 |

**Figure S12.** Total number of water molecules calculated from the simulated structure around the chromophore (*p*-HBI) in unbound state (gray), C1 bound state at 25 °C (salmon-red) and C1 bound state at 65 °C (orchid-purple). The result suggests that the [*p*-HBI]-loop-*Mj*RibK is more exposed to water in comparison to that at 25 °C.
